# GM-CSF-mediated epithelial-immune cell crosstalk orchestrates pulmonary immunity to *Aspergillus fumigatus*

**DOI:** 10.1101/2024.01.03.574062

**Authors:** Kathleen A. M. Mills, Frederike Westermann, Vanessa Espinosa, Eric Rosiek, Jigar V. Desai, Mariano A. Aufiero, Yahui Guo, Kennedy A. Mitchell, Selma Tuzlak, Donatella De Feo, Michail S. Lionakis, Amariliz Rivera, Burkhard Becher, Tobias M. Hohl

## Abstract

*Aspergillus fumigatus* causes life-threatening mold pneumonia in immune compromised patients, particularly in those with quantitative or qualitative defects in neutrophils. While innate immune cell crosstalk licenses neutrophil antifungal activity in the lung, the role of epithelial cells in this process is unknown. Here, we find that that surfactant protein C (SPC)-expressing lung epithelial cells integrate infection-induced IL-1 and type III interferon signaling to produce granulocyte-macrophage colony-stimulating factor (GM-CSF) preferentially at local sites of fungal infection and neutrophil influx. Using *in vivo* models that distinguish the role of GM-CSF during acute infection from its homeostatic function in alveolar macrophage survival and surfactant catabolism, we demonstrate that epithelial-derived GM-CSF increases the accumulation and fungicidal activity of GM-CSF-responsive neutrophils, with the latter being essential for host survival. Our findings establish SPC^+^ epithelial cells as a central player in regulating the quality and strength of neutrophil-dependent immunity against inhaled mold pathogens.

**HIGHLIGHTS:**

- GM-CSF is essential for host defense against *A. fumigatus* in the lung
- IL-1 and IFN-λ promote GM-CSF production by lung epithelial cells in parallel
- Epithelial cell-derived GM-CSF increases neutrophil accumulation and fungal killing capacity
- Epithelial cells preferentially upregulate GM-CSF in local sites of inflammation

**GRAPHICAL ABSTRACT:** 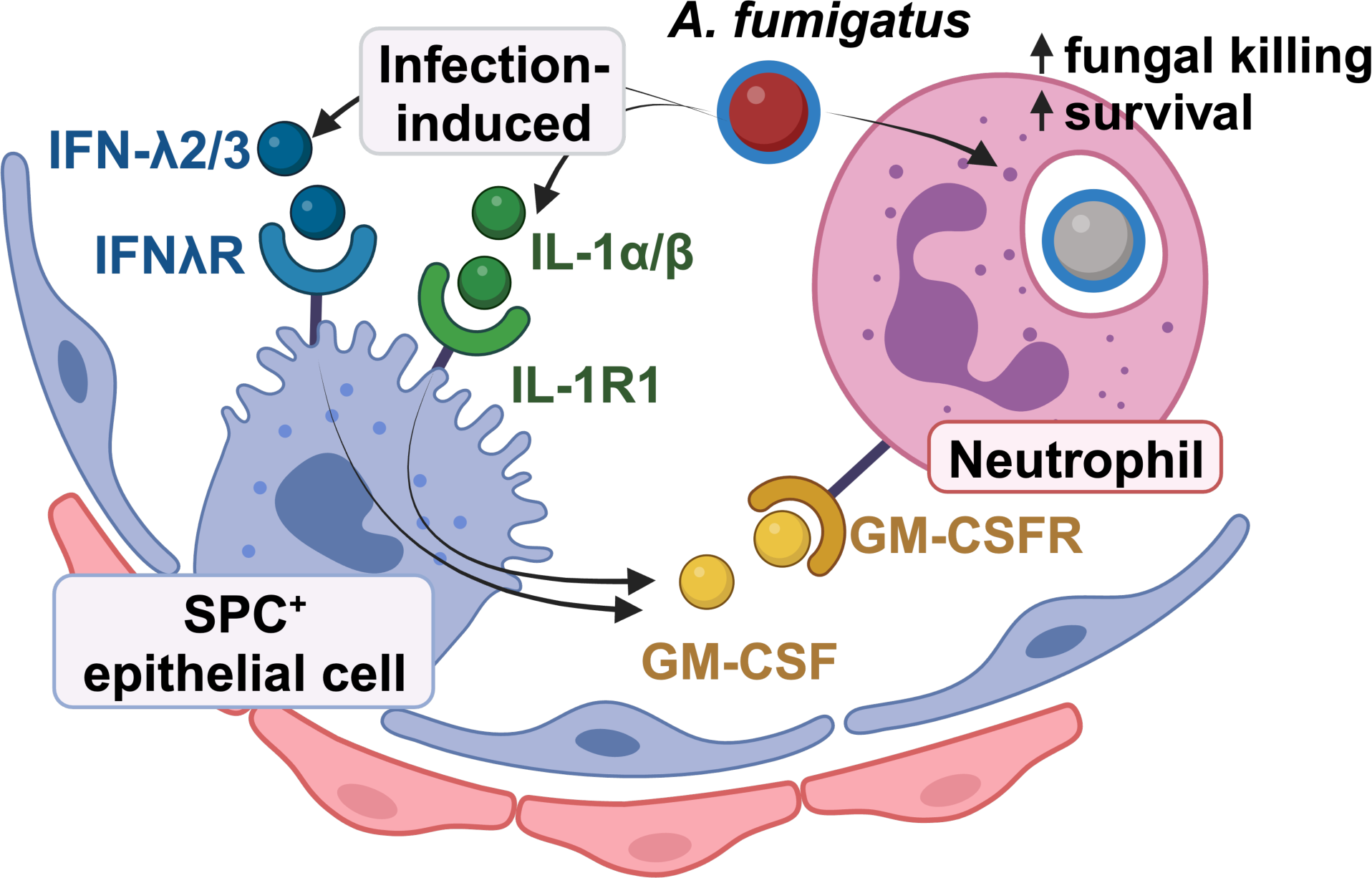

## INTRODUCTION

*Aspergillus fumigatus* is a ubiquitous opportunistic fungal pathogen that disseminates via airborne vegetative spores termed conidia.^1^ In immune compromised patients, conidia can germinate into tissue-invasive hyphae that have the capacity to invade the lung parenchyma and disseminate to remote sites. This process leads to the life-threatening disease invasive aspergillosis (IA). Myeloid cells are essential to maintain barrier immunity^2,3^, with prolonged neutropenia and monocytopenia as classic risk factors for IA. Qualitative defects in myeloid cell function (e.g., loss of neutrophil NADPH oxidase function in chronic granulomatous disease) also underlie susceptibility to IA.^1,4^

An emerging theme in antifungal immune defense is the recognition that recruited myeloid cells shape the pulmonary inflammatory environment and thereby license cellular antifungal effector functions. For example, CCR2^+^ monocyte- and neutrophil-derived interferons (IFNs) reciprocally enhance the oxidative burst^5^, and fungus-infected neutrophils and monocyte-derived dendritic cells (moDCs) release CXCR3 ligands that promote the infiltration of plasmacytoid dendritic cells (pDCs) that in turn enhance neutrophil fungicidal capacity. Our previous work has demonstrated that neutrophil killing of *A. fumigatus* conidia is increased by the presence of granulocyte-macrophage colony-stimulating factor (GM-CSF, encoded by *Csf2*).^6^ This is clinically relevant because patients with defects in GM-CSF signaling are susceptible to invasive fungal infection, including with *A. fumigatus*.^7^ However, these patients also develop pulmonary alveolar proteinosis (PAP) due to the loss of alveolar macrophages (AMs), which break down surfactant and pulmonary debris, a finding recapitulated in GM-CSF receptor (*Csf2rb*)^−/−^ mice.^8–12^ At homeostasis, AMs require GM-CSF production by type II alveolar epithelial cells (AECIIs) for their development and maintenance.^13^ While AMs can phagocytose and kill *A. fumigatus* in the infected lung, AMs are not essential for murine survival after infection^14^, suggesting that GM-CSF signaling mediates specific functions during infection that are distinct from those during the steady state.

These findings raise two distinct models of GM-CSF production and action. In the first model, GM-CSF production may be segregated by cell type to enact separate functions during tissue homeostasis and microbial infection. Support for this hypothesis comes from fungal infection models in which lymphocytes produce GM-CSF, exemplified by innate-like lymphocytes during pulmonary *Blastomyces dermatitidis* infection^15^ and NK cells during renal candidiasis.^16^ Additionally, during staphylococcal skin infection, γδ T cells make GM-CSF^17^, and in experimental autoimmune encephalomyelitis (EAE), CD4^+^ T cells are the primary source of GM-CSF in the central nervous system^18^. In the second model, GM-CSF may be derived from the same cell subsets under homeostatic and infectious conditions but require distinct signals for induction. *Rag2*^−/−^*Il2rg*^−/−^ mice that lack all lymphocytes have no defect in *A. fumigatus* clearance in the lung and do not develop IA,^19^ unlike *Csf2rb*^−/−^ mice, consistent with the idea that non-lymphoid cells may represent a critical source of GM-CSF during *A. fumigatus* infection. Further support for this idea comes from a murine model of pulmonary infection with the intracellular bacterial pathogen *Legionella pneumophila*, which triggered GM-CSF production by AECIIs, the same cell type that produces GM-CSF at homeostasis to mediate AM survival.^20^

In this study, we sought to understand the regulation, production, and function of GM-CSF in the *A. fumigatus-*infected lung, particularly by harnessing *in vivo* murine models that do not have PAP as a confounding factor. We demonstrate that IL-1 and type III IFN independently promote GM-CSF production early during *A. fumigatus* infection and that GM-CSF is principally derived from surfactant protein C (SPC)-expressing epithelial cells, including AECIIs.

Conditional deletion of the *Csf2* gene in SPC^+^ cells or of the *Csf2rb* gene in neutrophils results in defective neutrophil-mediated killing of *A. fumigatus* conidia and in increased murine mortality in the latter setting. SPC^+^ cells produce GM-CSF in proximity to *A. fumigatus* conidia and lung-infiltrating neutrophils, indicating that the local presence of inflammatory signals during fungal invasion activates epithelial cells to promote neutrophil antifungal defense at specific sites of infection. Our findings define an essential requirement for GM-CSF in host defense against *A. fumigatus* that is distinct from its homeostatic role in AM-mediated lung physiology. Further, they illuminate crosstalk between SPC^+^ epithelial cells and neutrophils to license neutrophil effector functions within spatially defined infectious foci.

## RESULTS

### *GM-CSF is required for host defense against* A. fumigatus

To determine the kinetics of GM-CSF production in the lung during *A. fumigatus* infection, we infected C57BL6/J (B6) mice intratracheally with 3 x 10^7^ resting conidia (CEA10 strain) and measured GM-CSF levels in lung homogenates by ELISA. GM-CSF production peaked at 12 hours post infection (hpi) and declined to baseline levels by 48 hpi (Fig. 1A, raw data from one experiment in Fig. S1A). To test whether GM-CSF is essential for survival of *A. fumigatus*-infected mice, we infected B6 and *Csf2*^−/−^ (GM-CSF knockout) mice and compared their mortality. 100% of *Csf2*^−/−^ mice died by 2 days post infection (dpi), while ∼30% of wild-type mice died by 4 dpi (Fig. 1B), indicating an essential role for GM-CSF during *A. fumigatus* infection. Since *Csf2*^−/−^ mice develop PAP, we hypothesized that death of these mice may be due to immunopathology caused by dysregulated inflammation and altered surfactant levels in the lung. To test this, we infected mice with swollen heat-killed conidia which expose immunostimulatory cell wall ligands (e.g., β-glucan and mannan epitopes) but are unable to germinate and form tissue-invasive hyphae. *Csf2*^−/−^ mice all survived infection with 3 x 10^7^ swollen heat-killed conidia (Fig. 1C), indicating that live *A. fumigatus* conidia are required for death of *Csf2*^−/−^ mice, and that exposure of immunostimulatory *A. fumigatus* cell wall components alone does not cause mortality in *Csf2^−/−^* mice.

**Fig. 1.**
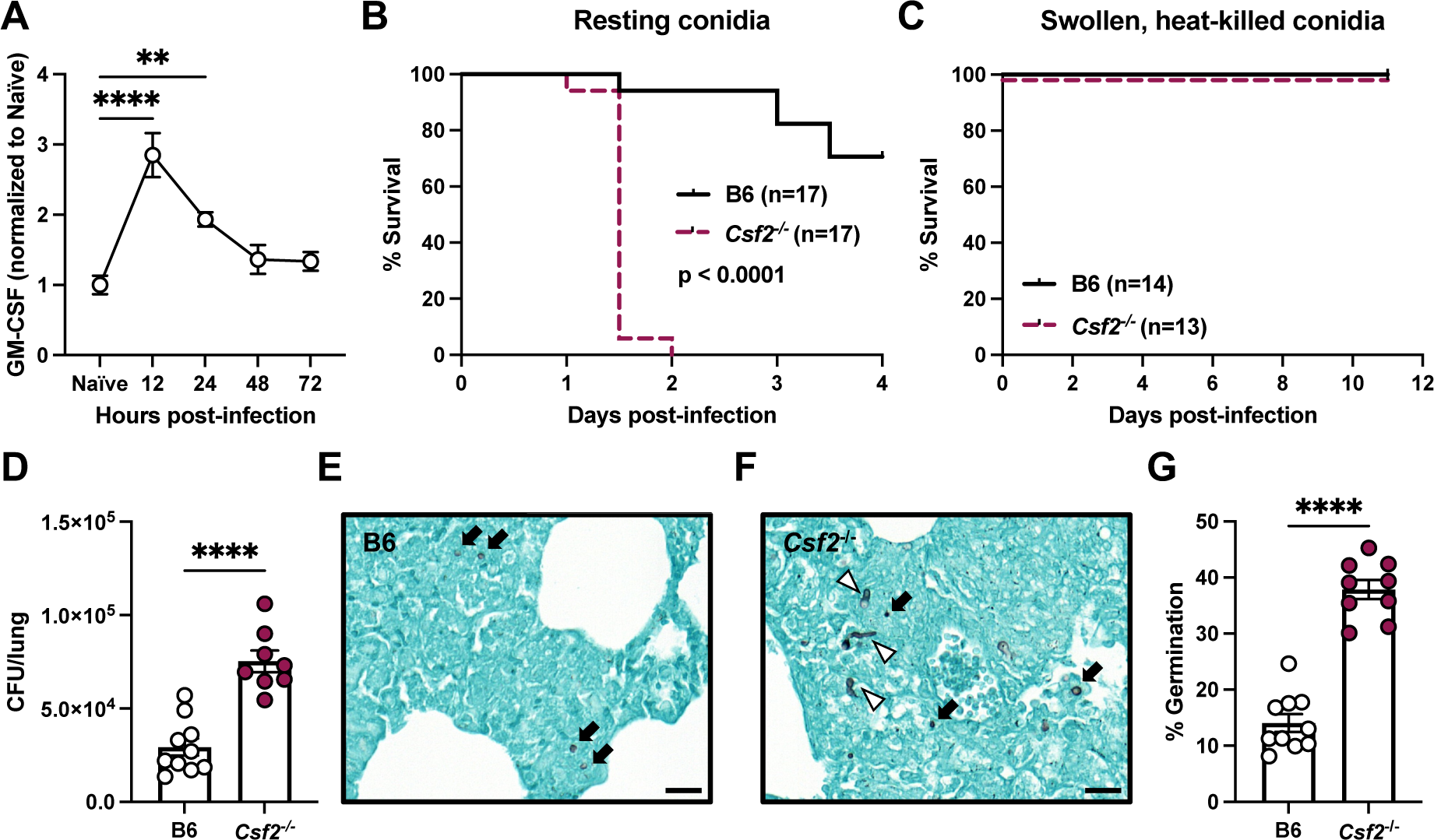
GM-CSF is required for host defense against *Aspergillus fumigatus*. (A) GM-CSF levels measured by ELISA in lung homogenates of naïve and *A.f.* infected (3 x 10^7^ conidia) B6 mice. (B) Survival of B6 and *Csf2*^−/−^ mice after infection with 4-6 x 10^7^ resting conidia. (C) Survival of B6 and *Csf2*^−/−^ mice after infection with 4-6 x 10^7^ swollen, heat-killed conidia. (D) CFU from lungs of B6 and *Csf2*^−/−^ mice 48 hpi with 1.5 x 10^7^ resting conidia. (E-F) Representative lung sections from B6 and *Csf2*^−/−^ mice infected with 1.5×10^7^ resting conidia 48 hpi stained with Grocott–Gomori’s methenamine silver (GMS). Black arrows indicate conidia and white arrowheads indicate germlings. Scale bar = 20 μm. (G) Percentage of germinating conidia quantified from GMS staining of lung sections from B6 and *Csf2*^−/−^ mice infected with 1.5×10^7^ resting conidia 48 hpi. Germinating conidia are defined as GMS^+^ regions with a maximum Feret diameter of >6μm. Each symbol represents one mouse. (A, D, G) were analyzed by Mann-Whitney test and are represented as mean ± SEM. (B & C) were analyzed by log-rank (Mantel-Cox) test. (A-D, G) are all pooled from 2 experiments. See also Fig. S1.

To investigate why *Csf2*^−/−^ mice are susceptible to *A. fumigatus*, we infected mice with a lower inoculum (1.5 x 10^7^ resting conidia) to allow for mice to survive to the experimental time point and quantified fungal burden in the lungs at 48 hpi by plating tissue homogenates and counting colony-forming units (CFU). *Csf2*^−/−^ mice had 2-3-fold higher lung CFUs compared to B6 control mice (Fig. 1D). We stained infected lungs from B6 and *Csf2*^−/−^ mice at 48 hpi with Gomori methenamine silver (GMS) stain to detect fungal organisms (Fig. 1E-1F). Using an observer-independent computational approach, we quantified the percentage of germinating conidia by calculating the maximum Feret diameter of all GMS-stained objects found in multiple annotated areas of GMS-stained slides (see Fig. S1B for a representative data analysis by annotation in one experiment). Since resting conidia are 2-3 μm in diameter and double in size upon swelling^21^, we designated any GMS^+^ object with diameter >6 μm as “germinating”. We found that *Csf2*^−/−^ mice had a significantly higher percentage of germinating conidia compared to B6 mice (Fig. 1G). Together, these data indicate that early production of GM-CSF is required during *A. fumigatus* infection to prevent mortality and restrict conidial germination into tissue-invasive hyphae.

### *IL-1 and type III IFN signaling promote GM-CSF production during* A. fumigatus *infection in parallel*

We sought to determine the upstream signaling pathway(s) that regulate GM-CSF production during *A. fumigatus* infection. We measured GM-CSF levels in the lungs of mice deficient in canonical fungal recognition pathways, including CARD9, Dectin-1, and FcRγ; the latter mediates signaling through multiple C-type lectin receptors, including Dectin-2 and Mincle. At 12 hpi, we found no difference in lung GM-CSF levels in *Card9*^−/−^, *Clec7a*^−/−^, or *Clec7a*^−/−^ *Fcer1g*^−/−^ mice relative to B6 controls (Fig. S2A-C). Thus, *A. fumigatus* ligands that stimulate Dectin-1 (β-glucans), FcRγ (mannans), and downstream CARD9 activation are not essential for rapid GM-CSF induction in the lung.

We investigated whether GM-CSF production during *A. fumigatus* infection may be downstream of interleukin (IL)-1 receptor signaling, as demonstrated in *Legionella pneumophila* and *Blastomyces dermatitidis* infection in the lung^15,20^ and in the EAE model of multiple sclerosis in the CNS^18^. Consistent with these observations, we found that at 12 hpi, GM-CSF levels in the lung were decreased in *Il1a/b*^−/−^ mice, *Il1r1*^−/−^ mice, and *Myd88*^−/−^ mice, which all lack components of the IL-1 signaling pathway (Fig. 2A-C). All mice with defects in IL-1 production or responsiveness contained approximately 50% lower GM-CSF levels in the lungs than control mice. Thus, we surmised that additional IL-1-independent signaling pathways may regulate GM-CSF production in the *A. fumigatus*-infected lung.

**Fig. 2.**
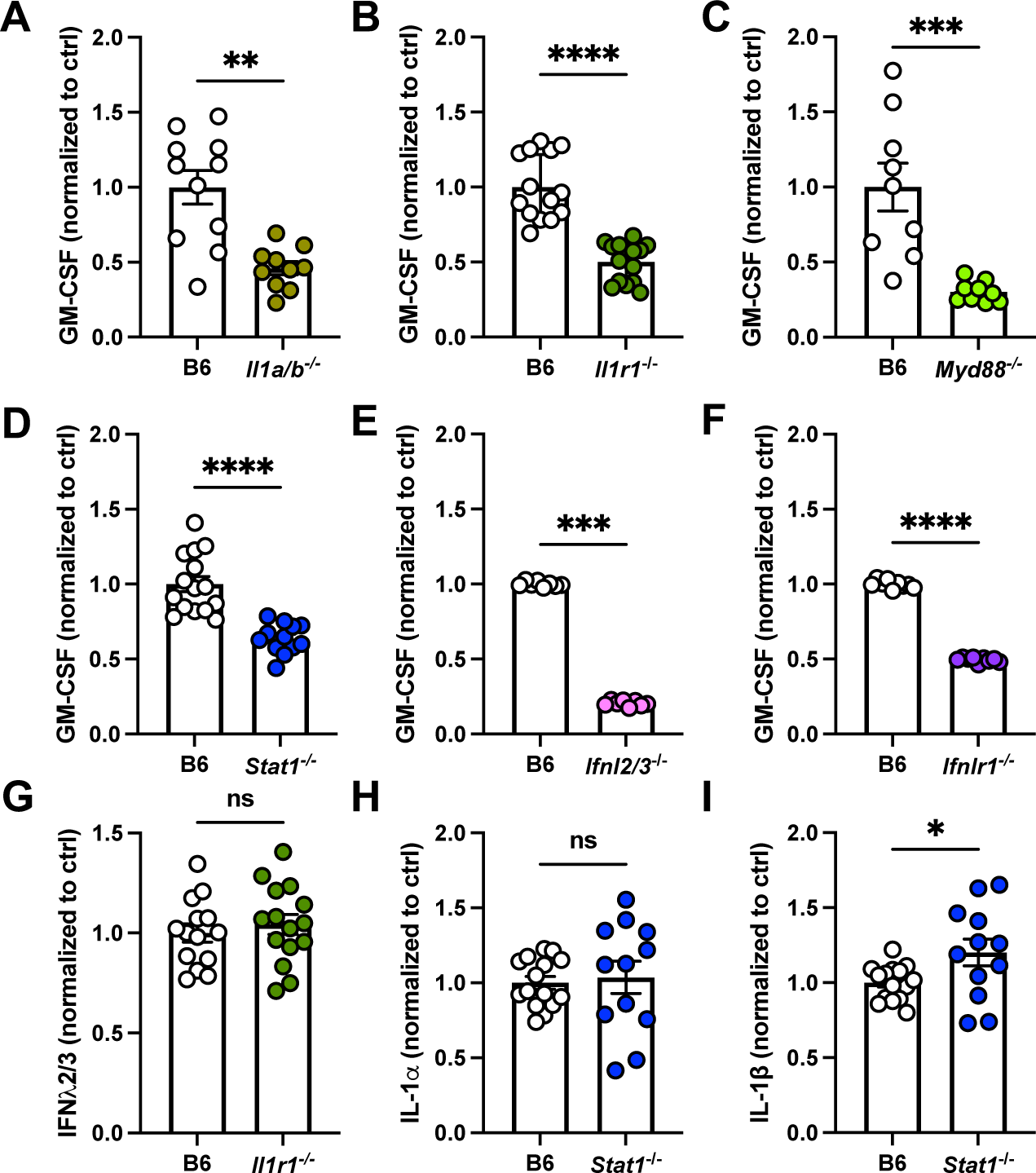
IL-1 and type III IFN signaling promote GM-CSF production during *A. fumigatus* infection in parallel. (A-F) GM-CSF levels measured by ELISA in B6 mice and (A) *Il1a/b*^−/−^, (B) *Il1r1*^−/−^, (C) *Myd88* ^−/−^, (D) *Stat1*^−/−^, (E) *Ifnl2/3*^−/−^, and (F) *Ifnlr1* ^−/−^ mice 12 hpi. (G) IFNλ2/3 levels measured by ELISA in B6 and *Il1r1*^−/−^ mice. (H) IL-1α and (I) IL-1β levels measured by ELISA in B6 and *Stat1*^−/−^ mice. (A-I) 2-3 experiments pooled; each symbol represents one mouse and data is displayed as mean ± SEM. Significance determined by Mann-Whitney test. See also Fig. S2.

Type I (α and β) and III (λ) interferons (IFN), classically thought of as anti-viral cytokines, have been recently described to have important roles in defense against pulmonary *A. fumigatus* infection and are produced at early stages post-infection.^19^ To determine whether signaling through IFN receptors regulates GM-CSF production during *A. fumigatus* infection, we quantified GM-CSF in the lungs of *Stat1*^−/−^ mice, which lack the Stat1 transcription factor downstream of type I and III IFN pathways^22^. We found that *Stat1*^−/−^ mice had impaired GM-CSF production compared to B6 controls, consistent with a contribution of IFN receptor signaling to GM-CSF production (Fig. 2D). To determine which IFNs promote GM-CSF production, we infected mice deficient in the type I IFN receptor, IFNAR1, and mice deficient in the type III IFN cytokines IFN-λ2/3 and their receptor IFNLR1. We did not focus on IFN-γ (type II IFN) signaling because a prior study demonstrated that *Ifngr1*^−/−^ mice were not susceptible to *A. fumigatus* challenge, unlike *Ifnar1*^−/−^ and *Ifnlr1*^−/−^ mice.^19^ *Ifnar1*^−/−^ mice had normal levels of GM-CSF at 12 hpi (Fig. S2D), but both *Ifnl2/3*^−/−^ mice and *Ifnlr1*^−/−^ mice had reduced GM-CSF in the lungs at that time point (Fig. 2E-2F). These results demonstrate that type III, but not type I, IFN signaling promotes GM-CSF production during *A. fumigatus* infection.

While lung GM-CSF production relied on both IL-1 and IFN-λ signaling, it remained unclear whether these pathways operate in parallel or sequentially in the context of *A. fumigatus* infection. To test whether IFN-λ production is downstream of IL-1 signaling, we measured IFN-λ2/3 levels in the lungs of infected WT and *Il1r1*^−/−^ mice. We found no difference in IFN-λ2/3 levels between these groups (Fig. 2G). To test whether IL-1 production is downstream of IFN-λ signaling, we measured IL-1α and IL-1β levels in the lungs of infected WT and *Stat1*^−/−^ mice and found no difference between these groups (Fig. 2H-2I). Taken together, these results demonstrate that the IL-1-dependent and IFN-λ-dependent GM-CSF production are independent and promote GM-CSF production in parallel.

### IL-1 and Stat1 are individually dispensable for GM-CSF production at steady state

While GM-CSF production by AECIIs is known to be essential for homeostatic development and function of alveolar macrophages^13^, the pathways that control GM-CSF production at steady state are unknown. We hypothesized that basal IL-1 or IFN-λ production may promote GM-CSF production at homeostasis, beyond their regulatory role during infection. To investigate this idea, we quantified alveolar macrophages in the lungs of uninfected *Il1r1*^−/−^ and *Stat1*^−/−^ mice. We found that alveolar macrophage numbers in both groups of gene-deficient mice were unchanged compared to B6 mice (Fig. 3A-B). In addition, we measured GM-CSF levels by ELISA in the lungs of uninfected mice and found no difference between B6, *Il1r1*^−/−^, and *Stat1*^−/−^ mice (Fig. 3C). These results indicate that while IL-1 and Stat1 control GM-CSF production during *A. fumigatus* infection, these pathways are individually dispensable for GM-CSF production at homeostasis.

**Fig. 3.**
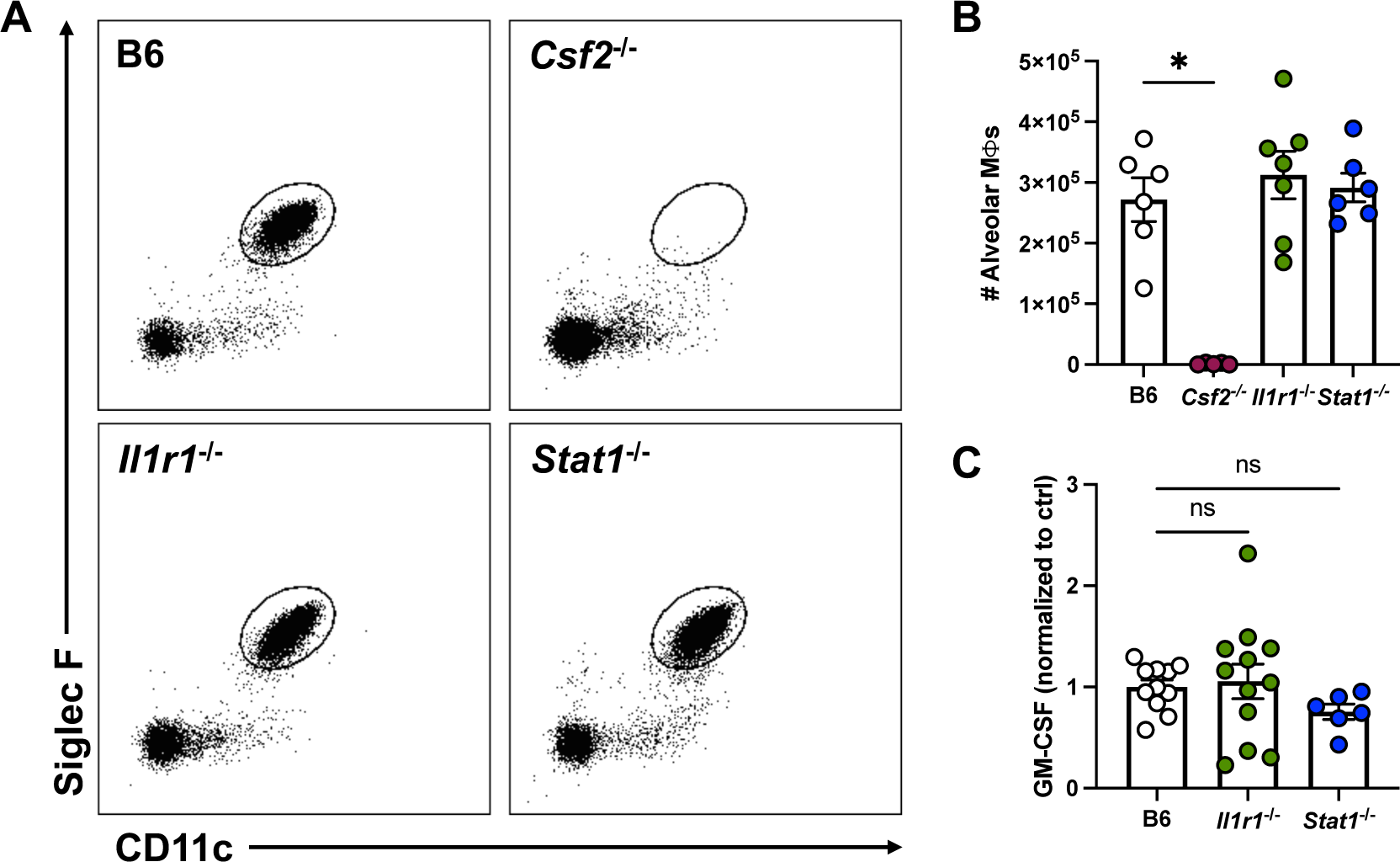
IL-1 and Stat1 are individually dispensable for GM-CSF production at homeostasis. (A) Representative flow plots from lungs of uninfected B6, *Csf2*^−/−^, *Il1r1*^−/−^, and *Stat1*^−/−^ mice, gated on CD45^+^ live cells. Black gate indicates alveolar macrophages (CD11c^+^ Siglec F^+^). (B) Number of alveolar macrophages and (C) GM-CSF levels measured by ELISA in the lungs of uninfected B6, *Csf2*^−/−^, *Il1r1*^−/−^, and *Stat1*^−/−^ mice. (B & C) 2-3 experiments pooled. Each symbol represents one mouse and data is displayed as mean ± SEM. Significance determined by Kruskal-Wallis test.

### GM-CSF is produced by Surfactant Protein C^+^ pulmonary epithelial cells

To determine which cell compartment produces GM-CSF during *A. fumigatus* infection, we generated bone marrow chimeras by irradiating recipient WT (CD45.1^+^) mice or *Csf2*^−/−^ (CD45.2^+^) mice and reconstituting them with donor WT or *Csf2*^−/−^ bone marrow cells. Six weeks post reconstitution, we infected them with *A. fumigatus* conidia. At 12 hpi, the peak of GM-CSF production (Fig. 1A), lung GM-CSF levels were reduced in both groups of mice with *Csf2*-deficient radioresistant cells, but not in mice with *Csf2*-deficient radiosensitive cells (Fig. 4A), indicating that radioresistant cells are the primary source of GM-CSF during infection. To determine whether radioresistant cell-derived GM-CSF is important for murine survival, we infected chimeras and found that mice with *Csf2*-deficient radioresistant cells had reduced survival compared to mice with *Csf2*-deficient radiosensitive cells (Fig. 4B). Thus, GM-CSF produced by pulmonary radioresistant cells is required for murine survival following *A. fumigatus* infection.

**Fig. 4.**
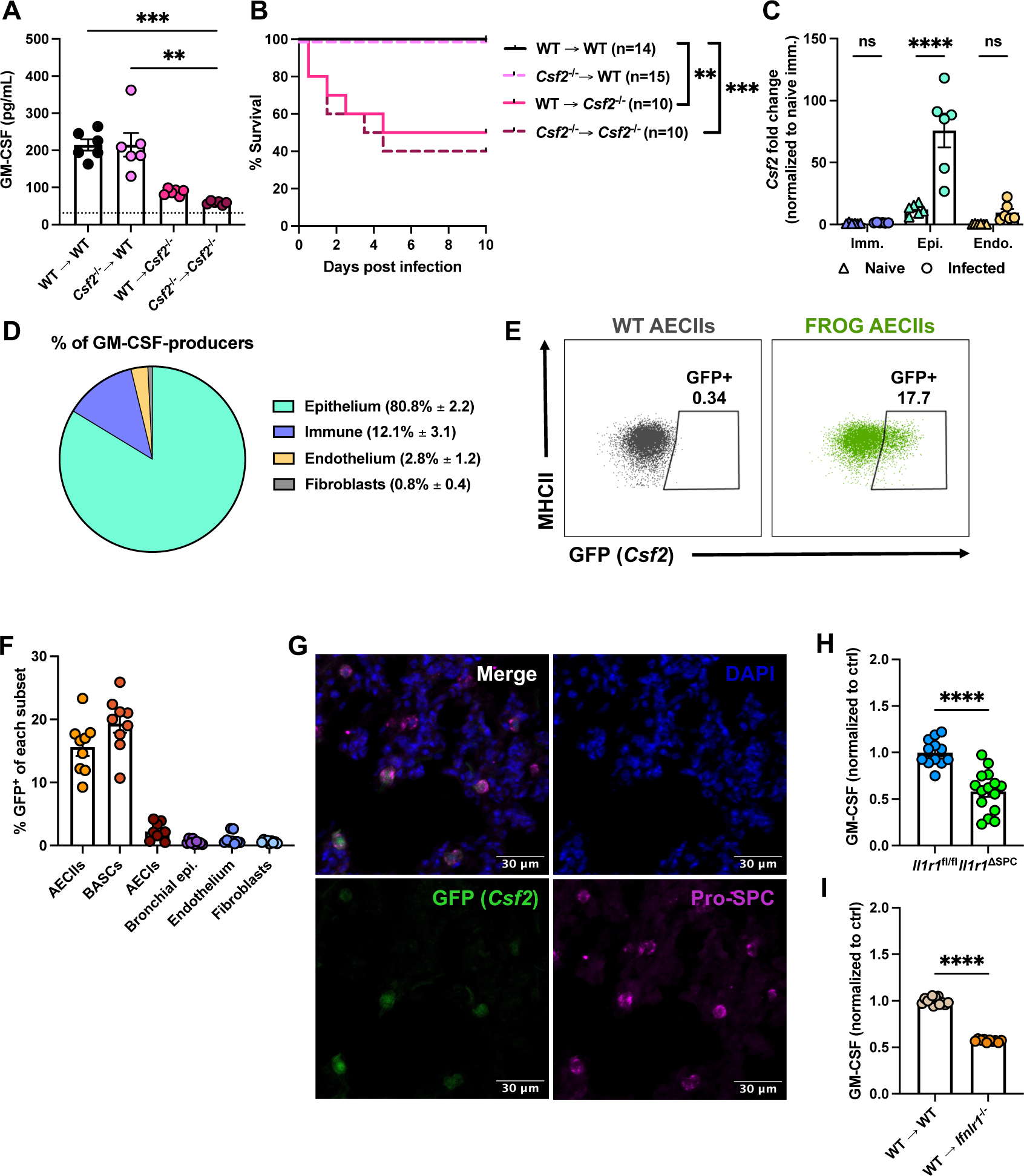
GM-CSF is produced by Surfactant Protein C^+^ pulmonary epithelial cells. (A) GM-CSF levels measured by ELISA in *Csf2*^−/−^ bone marrow chimeras 12 hpi. Dotted line represents lower limit of detection of the assay. (B) Survival of *Csf2*^−/−^ chimeras after *A.f.* infection. 2 experiments pooled (significance calculated by log-rank test). (C) *Csf2* transcript levels measured by qPCR in FACS sorted cells (see Fig. S2 for gating strategy) from naïve or infected B6 mice. 2 experiments pooled; each symbol is one mouse. Data normalized to naïve immune cells; bars represent mean ± SEM. (D) Pie chart displaying fraction of GFP (*Csf2*) positive cells after *A.f.* infection. Summary data from four mice (2 experiments) are displayed as mean ± SD. (E) Representative flow plots of AECIIs from WT and FROG mice displaying GFP^+^ cells. (F) Percent of GFP (*Csf2*) positive cells out of each indicated subset as measured by flow cytometry in *A.f.* infected mice. Data are from 3 pooled experiments; each symbol is one mouse. (G) Immunofluorescence images of infected FROG mouse lung. (H) GM-CSF levels measured by ELISA in *Il1r1*^fl/fl^ and *Il1r1*^ΔSPC^ mice at 12 hpi. Data are from 3 pooled experiments. (I) GM-CSF levels measured by ELISA in irradiated WT or *Ifnlr1*^−/−^ recipient mice reconstituted with WT bone marrow at 12 hpi. Data are from 2 pooled experiments. (A, C, F, H & I) Each symbol represents one mouse. (A) is analyzed by Kruskal-Wallis test. (B) is analyzed by log-rank (Mantel-Cox) test. (C) is analyzed by two-way ANOVA. (H & I) are analyzed by Mann-Whitney test. See also Fig. S3.

To define which radioresistant cell types express *Csf2* during infection, we sorted CD45^+^ immune, CD45^−^ EpCAM^+^ epithelial, and CD45^−^ CD31^+^ endothelial cells by FACS (gating strategy in Fig. S3A) from the lungs of naïve and infected B6 mice at 6 hpi and performed qPCR for *Csf2* on each cell subset. Epithelial cells had the highest expression of *Csf2*, both at steady state and during infection, with a significant increase in expression between those states, while endothelial cells and immune cells had limited *Csf2* expression at steady state and during infection (Fig. 4C).

To investigate further which epithelial cell subsets make GM-CSF during infection, we infected fluorescent reporter <u>o</u>f <u>G</u>M-CSF (FROG) mice^18^, a transgenic reporter mouse line with GFP inserted in the native *Csf2* locus, and examined GFP expression in single cell lung suspensions by flow cytometry. By gating first on live GFP^+^ cells, we found that ∼80% of *Csf2*-expressing cells were epithelial cells (CD45^−^ EpCAM^+^), with minor contributions from the immune cell compartment (CD45^+^ cells; Fig. 4D). Within the epithelial cell compartment, *Csf2* expression was confined to type II alveolar epithelial cells (AECIIs) and to bronchoalveolar stem cells (BASCs) and was absent in type I alveolar epithelial cells (AECIs) and in bronchial epithelial cells (Fig. 4E-4F). The identity of AECIIs and BASCs was confirmed by sorting these cells from uninfected mouse lungs and by quantifying *Sftpc* (encoding surfactant protein C [SPC]), *Scgb1a1*, and *Csf2* expression (Fig. S3B-S3D), compared to sorted endothelial cells as a negative control. AECIIs and BASCs both express *Sftpc*, and BASCs additionally express *Scgb1a1*, while AECIIs do not.^23,24^ Therefore, we refer to AECIIs and BASCs as “SPC^+^ cells” for the remainder of the manuscript. Within the immune cell compartment, GFP expression was restricted to γδ Τ cells and ILCs, and not found in other lymphoid or myeloid cell subsets (Fig. S3E-S3L). Finally, endothelial cells and fibroblasts had minimal to no expression of *Csf2* (Fig. 4D-4F).

To confirm further that SPC^+^ cells were GFP^+^, we infected FROG mice with *A. fumigatus*, and examined pro-SPC and GFP expression in the lungs by immunofluorescence and confocal microscopy. We found that GFP^+^ cells overlapped with pro-SPC^+^ cells, but not all pro-SPC^+^ cells were GFP^+^ (Fig. 4G). This finding was consistent with the flow cytometry data that indicated that 15-20% of SPC^+^ cells expressed GFP (Fig. 4F).

To determine whether IL-1R1 signaling is required on SPC^+^ epithelial cells for GM-CSF production, we deleted *Il1r1* in SPC^+^ cells by crossing *Il1r1*-floxed mice to *Sftpc*-Cre^ERT2^ mice. After tamoxifen administration, efficient deletion of *Il1r1* was achieved in both AECIIs and BASCs as measured by qPCR in cells sorted from uninfected mice (Fig. S3M-S3N). We found that *Il1r1*^ΔSPC^ mice had reduced lung GM-CSF levels compared to Cre-negative *Il1r1*^fl/fl^ littermate controls (Fig. 4H).

To determine whether IFN-λ signaling is required on radioresistant cells for GM-CSF induction, we generated bone marrow chimeras using WT bone marrow and either WT or *Ifnlr1*^−/−^ irradiated recipient mice. At 12 hpi with *A. fumigatus*, GM-CSF was reduced in the *Ifnlr1*^−/−^ recipient group, indicating a requirement for IFN-λ signaling on radioresistant cells for optimal GM-CSF production (Fig. 4I). Taken together, these data indicate that SPC^+^ pulmonary epithelial cells are the relevant producers of GM-CSF during *A. fumigatus* infection and respond both to IL-1 and to IFN-λ signaling to mediate infection-induced GM-CSF production.

### GM-CSF-producing SPC^+^ cells form a signaling circuit in proximity to A. fumigatus and lung-infiltrating neutrophils

We hypothesized that heterogeneity in GM-CSF production by SPC^+^ epithelial cells (∼15-20% of these cells are *Csf2*^+^ during infection by flow cytometric quantitation of GFP expression; Fig. 4F) may be explained by their proximity to *A. fumigatus* or neutrophils at local sites of infection (Fig. 5A). To test this idea, we quantified the minimum distance from GFP^+^ pro-SPC^+^ or GFP^−^ pro-SPC^+^ cells to AF633-labeled *A. fumigatus* by fluorescence microscopy using scans of whole lobes of lung tissue from infected FROG mice (Fig. 5B). We found that GFP^+^ pro-SPC^+^ cells were in closer proximity to *A. fumigatus* compared to GFP^−^ pro-SPC^+^ cells, consistent with the model that the local presence of *A. fumigatus* facilitates *Csf2* upregulation by SPC^+^ epithelial cells via IL-1 and type III IFN dependent pathways (Fig. 5C, see breakdown by mouse in Fig. S4A-G).

**Fig. 5.**
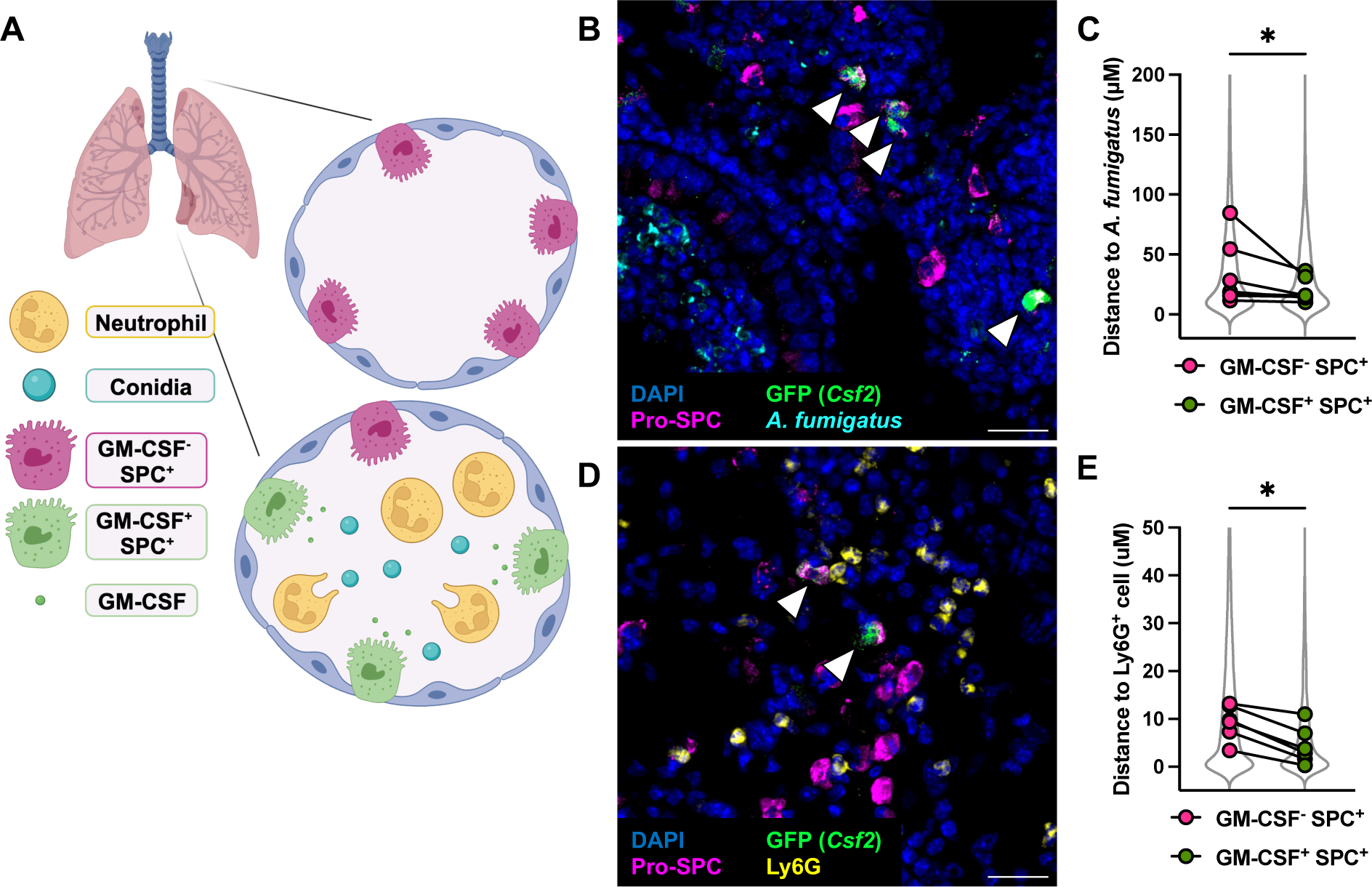
GM-CSF-producing epithelial cells form a signaling circuit in proximity to *A. fumigatus* and neutrophils. (A) Schematic of hypothesis that GM-CSF-producing SPC^+^ cells are in closer proximity to conidia and neutrophils than GM-CSF-non-producing SPC^+^ cells. (B) Representative image of mouse lung 24 hpi with AF633^+^ conidia stained with DAPI, anti-GFP, and pro-SPC. (C) Distance from indicated cell type to AF633^+^ conidia. Each dot is the median of distances for each indicated cell type, paired values refer to samples from one mouse (n = 3 mice). Dots are superimposed on violin plots representing all cells analyzed in a SuperPlot.^54^ (D) Representative image of mouse lung 24 hpi stained with DAPI, anti-GFP, pro-SPC, and Ly6G. (E) Distance from indicated cell type to Ly6G^+^ cell. Each dot is the median of distances for each indicated cell type, paired values refer to samples from one mouse (n = 5 mice). Dots are superimposed on violin plots representing all cells analyzed. See also Fig. S4. (B & D) White arrowheads indicate GM-CSF^+^ pro-SPC^+^ cells. Scale bar = 20 μM. (C & E) analyzed by Wilcoxon test.

In addition, we measured the minimum distance from either GFP^+^ pro-SPC^+^ or GFP^−^ pro-SPC^+^ cells to Ly6G^+^ cells (a canonical neutrophil marker; Fig. 5D). We found that GFP^+^ pro-SPC^+^ cells were in closer proximity to Ly6G^+^ cells compared to GFP^−^ pro-SPC^+^ cells (Fig. 5E, see breakdown by mouse in Fig. S4H-M), indicating that GM-CSF-producing epithelial cells are in spatial proximity to downstream cellular targets. Together, these data indicate that GM-CSF-producing SPC^+^ epithelial cells form a regional network of cell-cell communication with *A. fumigatus* and Ly6G^+^ neutrophils.

### GM-CSF derived from SPC^+^ epithelial cells is required for neutrophil fungicidal activity

Since GM-CSF is primarily produced by SPC^+^ epithelial cells, our model would predict that deleting *Csf2* in these cells would reduce the total amount of GM-CSF in the lung and have adverse effects on the antifungal properties of GM-CSF-responsive myeloid effector cells. To test this idea, we generated mice in which *Csf2* is deleted by tamoxifen administration in *Sftpc*-expressing cells by crossing *Csf2*-floxed (first described in this study; Fig. S5A) and *Sftpc*-Cre^ERT2^ mice. Importantly, in contrast to *Csf2*^−/−^ mice, these mice do not have a developmental defect in alveolar macrophages. We infected *Csf2*^ΔSPC^ or control mice with *A. fumigatus*, and measured GM-CSF in the lungs at 12 hpi. Deleting *Csf2* in SPC^+^ cells reduced the total amount of GM-CSF in the lungs (Fig. 6A), confirming that SPC^+^ cells are the primary producers of GM-CSF during *A. fumigatus* infection. Additionally, *Csf2*^ΔSPC^ mice had a defect in neutrophil accumulation in the lungs at 24 hpi compared to controls (Fig. 6B, gating strategy in Fig. S5B). Alveolar macrophage numbers were also reduced by ∼90% in *Csf2*^ΔSPC^ mice two weeks after tamoxifen administration, consistent with prior work^13^ (Fig. S5C).

**Fig. 6.**
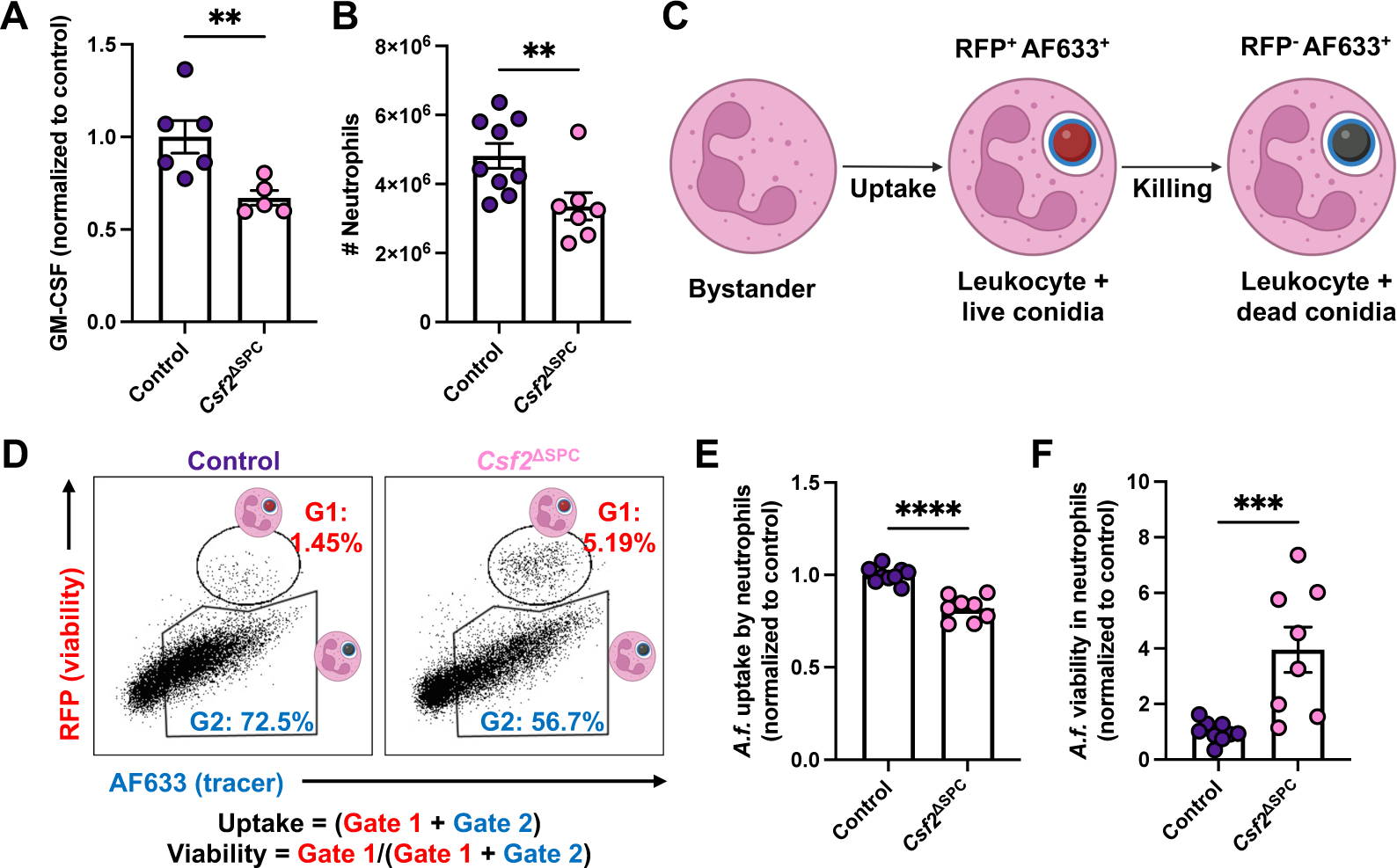
GM-CSF derived from SPC^+^ epithelial cells is required for neutrophil fungicidal activity. (A) GM-CSF levels measured by ELISA in *Csf2*^Δ*SPC*^ mice treated with vehicle or tamoxifen 12 hpi. (B) Neutrophil numbers in lungs of control and *Csf2*^Δ*SPC*^ mice 24 hpi. (C-D) Schematic of Fluorescent *Aspergillus* Reporter (FLARE) conidia and fluorescence emission after uptake and killing by leukocytes. (E) Uptake of conidia by and (F) conidial viability in lung neutrophils, quantified using FLARE conidia and flow cytometry from infected control and *Csf2*^Δ*SPC*^ mice 24 hpi. (A, B, E & F) Data are from two pooled experiments. Each symbol represents one mouse. Data were analyzed by Mann-Whitney test. See also Fig. S5.

To investigate whether *Csf2* expression by SPC^+^ epithelial cells is required for neutrophil fungal uptake or killing, we infected *Csf2*^ΔSPC^ or control mice with <u>fl</u>uorescent *<u>A</u>spergillus* <u>re</u>porter (FLARE) conidia^25^ (Fig. 6C-D). FLARE conidia encode a constitutively expressed RFP and are surface labeled with AF633. Upon phagocytosis of live FLARE conidia, host leukocytes emit RFP and AF633 fluorescence, but upon fungal killing, RFP is rapidly degraded in the phagolysosome while AF633 fluorescence is maintained, resulting in AF633^+^RFP^−^ leukocytes. Leukocyte fungal uptake is quantified as the fraction of a given leukocyte population that is AF633^+^. To quantify *Aspergillus* viability within leukocytes that have engulfed conidia, the frequency of RFP^+^ leukocytes is divided by the frequency of all AF633^+^ leukocytes (Fig. 6D). By flow cytometry, we found that neutrophils in the lungs of infected *Csf2*^Δ*SPC*^ mice had a defect in the uptake of *A. fumigatus* conidia (Fig. 6E) and contained more live *A. fumigatus* conidia compared to neutrophils in littermate controls (Cre-negative *Csf2*^fl/fl^; Fig. 6F). These data indicate that GM-CSF derived from SPC^+^ epithelial cells is required for neutrophils to accumulate in the infected lung and license their full fungicidal capability.

### GM-CSF signaling on neutrophils is required for conidial killing and murine survival

To determine whether GM-CSF receptor signaling on neutrophils is essential for their cell-intrinsic fungicidal activity and host defense in the lung, we deleted *Csf2rb* (encoding the β chain of the GM-CSF receptor) in neutrophils by crossing *Csf2rb*-floxed mice^26^ to *Mrp8*-cre mice (*Csf2rb*^ΔNeuts^). Deletion was confirmed by sorting lung neutrophils from infected mice and quantifying *Csf2rb* expression by qPCR (Fig. S6). Importantly, *Csf2rb*^ΔNeuts^ mice do not develop PAP since alveolar macrophages normally express the GM-CSF receptor. Compared to littermate control (Cre-negative *Csf2rb*^fl/fl^) mice, *Csf2rb*^ΔNeuts^ mice had reduced neutrophil accumulation into the lung (Fig. 7A), akin to the results observed in *Csf2*^ΔSPC^ mice. To investigate whether *Csf2rb* expression on neutrophils was required for fungal uptake or killing, we infected *Csf2rb*^ΔNeuts^ mice with FLARE conidia (Fig. 7B). Neutrophils in the lungs of *Csf2rb*^ΔNeuts^ mice had similar levels of fungal uptake compared to littermate controls but were more likely to harbor viable *A. fumigatus* conidia, indicating that GM-CSF signaling on neutrophils is dispensable for fungal uptake but required for optimal fungal killing (Fig. 7C & D). Finally, following *A. fumigatus* challenge, *Csf2rb*^ΔNeuts^ mice had decreased survival compared to littermate control mice (Fig. 7E). Taken together, these results indicate that GM-CSF signaling on neutrophils protects mice from mortality by promoting neutrophil accumulation and *A. fumigatus* conidial killing.

**Fig. 7.**
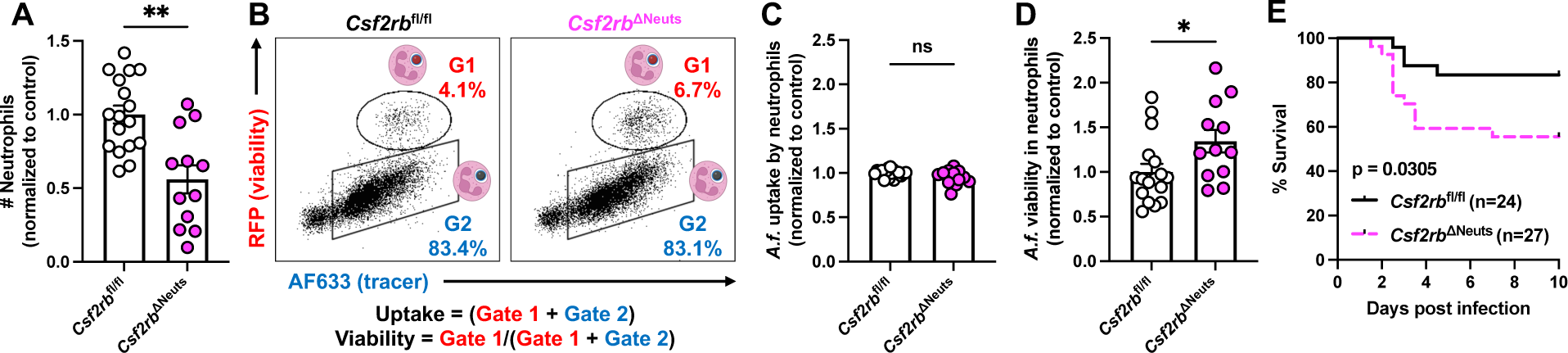
GM-CSF signaling on neutrophils is required for conidial killing and murine survival. (A) Neutrophil numbers in lungs of *Csf2rb*^fl/fl^ and *Csf2rb*^ΔNeuts^ mice 24 hpi quantified by flow cytometry. (B) Representative dot plots gated on neutrophils in *Csf2rb*^fl/fl^ and *Csf2rb*^ΔNeuts^ mice 24 hpi. (C) Uptake of conidia by and (D) conidial viability in lung neutrophils 24 hpi. (D) Survival of *Csf2rb*^fl/fl^ and *Csf2rb*^ΔNeuts^ mice after infection with 4-6×10^7^ *A.f.* conidia. 4 pooled experiments. (A, C-D) Multiple experiments pooled, significance calculated by Mann-Whitney test. (E) analyzed by log-rank (Mantel-Cox) test. See also Fig. S6.

## DISCUSSION

In this study, we sought to describe the role of GM-CSF in immune defense against an inhaled mold pathogen. In doing so, we have identified a critically important relationship between epithelial cells and neutrophils in invasive pulmonary fungal disease. We found that SPC^+^ pulmonary epithelial cells produce GM-CSF downstream of IL-1 and IFN-λ signaling to promote neutrophil killing of *A. fumigatus* conidia in spatially distinct foci of infection. These findings suggest that the susceptibility of PAP patients to invasive aspergillosis is attributable to loss of acute lung epithelial-derived GM-CSF that regulates neutrophil antifungal activity. Although GM-CSF originates from the same epithelial cell subset during homeostasis to regulate AM survival and surfactant catabolism, during acute mold pneumonia, infection-induced IL-1- and type III IFN-dependent regulation enable precise calibration of GM-CSF levels under these distinct conditions.

An emerging concept is that recruited innate immune cells are licensed at the site of infection to achieve their full antifungal capacity. Previous work by our group has demonstrated that pDCs license neutrophils to boost their fungicidal capacity at late time points (72 hpi) during *A. fumigatus* infection, increasing ROS production by those cells.^3^ In mice depleted of pDCs, neutrophil recruitment to the lung was intact, but neutrophil killing of conidia was decreased, indicating that pDC-derived signals are required for local instruction of neutrophils. Similarly, Espinosa et al. found that depletion of neutrophils has no impact on monocyte recruitment to the *A. fumigatus-*infected lung, but that monocytes from neutrophil-depleted mice are less mature, exhibit lower ROS production, and have diminished fungicidal activity (at 36 hpi).^5^ Conversely, neutrophils from monocyte-depleted mice are also less fungicidal (at 36 hpi).^2^ These data suggest that multiple immune cell subsets collaborate to promote fungicidal activity in the infected lung. Outside the lung, T cell-derived GM-CSF licenses mature monocytes to produce ROS in the CNS during EAE.^27^ Our findings in this study expand this crosstalk concept that was previously restricted to immune cells to incorporate epithelial cells. We found that epithelial cells respond to the upstream cytokine signals of IL-1 and IFN-λ to license neutrophils for optimal fungal uptake and killing via GM-CSF signaling. Additionally, our data indicate that this process of licensing occurs at the earliest stages of infection with the cells that are present in the lung at that time, i.e., epithelial cells and neutrophils (12-24 hpi). Thus, GM-CSF-dependent epithelial-neutrophil crosstalk occurs before bidirectional monocyte-neutrophil crosstalk (peak at 36 hpi) and pDC-neutrophil crosstalk (peak at 72 hpi).

Prior to this study, the cellular source(s) of GM-CSF during *A. fumigatus* infection were unknown. In other models of fungal infection, including *Blastomyces dermatitidis* in the lung^15^ and *Candida albicans* in the kidney^16^, innate lymphocytes are an important source of GM-CSF. Additionally, in the steady-state intestine, ILCs are a major source of GM-CSF.^28^ While lymphocytes did contribute ∼15% of the transcriptional pool of *Csf2* in our model (Fig. S3E-S3F), deleting *Csf2* in SPC^+^ cells was sufficient to impair neutrophil fungicidal activity, indicating that lymphocyte-derived GM-CSF cannot compensate for epithelial-cell derived GM-CSF during *A. fumigatus* infection. Whether non-immune cells are relevant sources of GM-CSF in other models of fungal infection remains unexplored. However, during pulmonary bacterial infection with a *Legionella pneumophila* mutant, SPC^+^ epithelial cells (AECIIs) represent a critical source of GM-CSF.^20^ Additionally, air-liquid interface cultures of murine tracheal and human bronchial epithelial cells produce GM-CSF after allergenic challenge with house dust mite extract.^29^

Our study provides novel insights into the upstream regulation of GM-CSF during *A. fumigatus* infection. Here, we demonstrate that GM-CSF production is co-regulated by IL-1 and IFN-λ. IL-1-dependent regulation of GM-CSF has been previously observed in other systems, including in the CNS during autoimmunity^18^, in the intestine at steady-state^28^, during pulmonary bacterial infection^20^, and during allergenic challenge in the lung^29^. Prior work from our group had identified IL-1R1 and MyD88 signaling on radio-resistant cells as important for anti-*A. fumigatus* immunity, particularly for neutrophil accumulation in the lung.^30^ Here, we found that loss of *Csf2* in SPC^+^ epithelial cells or loss of *Csf2rb* in neutrophils also impaired neutrophil accumulation in the lung, indicating that IL-1-dependent, epithelial-derived GM-CSF has a non-redundant role in neutrophil migration or turnover. Future studies may explore the cellular sources of IL-1R1 ligands and whether IL-1α or IL-1β are required for GM-CSF production in this setting. Prior work suggests a greater role for IL-1α than IL-1β in anti-*Aspergillus* immunity in otherwise immune competent mice.^31,32^ In contrast, the NLRP3 and AIM2 inflammasomes are linked to IL-1β production following *A. fumigatus* challenge and play an important role in host defense in corticosteroid-treated mice.^33,34^

Unlike IL-1, links between IFN-λ and GM-CSF are few, although *Ifnlr1*^−/−^ mice have a reduction in GM-CSF-producing T cells in the CNS during EAE.^35^ In this study, we report a direct signaling axis in which infection-induced IFN-λ responsiveness regulates epithelial-derived GM-CSF in the lung. Recent work has demonstrated a critical role for IFN-λ during *A. fumigatus* infection, particularly at late time points (48-72 hpi).^19^ At these time points, conditional deletion of IFNLR1 on neutrophils results in a reduction in neutrophil ROS production and elevated fungal CFU in the lungs. Bone marrow chimeras generated using *Ifnlr1*^−/−^ mice indicate a requirement for IFNLR1 expression on radiosensitive cells for murine survival. While loss of IFNLR1 expression on radioresistant cells was not required for murine survival in that study, IFNLR1 is known to be expressed by the lung epithelium and epithelial cells are an important mediator of IFNLR1 signaling during viral infection.^36,37^

Here, we found that IFNLR1 expression on radioresistant cells is required for GM-CSF production during *A. fumigatus* infection. However, in this bone marrow chimera setting, IL-1-dependent GM-CSF production likely compensates at least in part for IFN-λ-dependent GM-CSF production to protect mice against mortality. These data indicate that IL-1R1 and IFNLR1 expression by epithelial cells is redundant for a certain threshold of GM-CSF production, providing a beneficial outcome for organismal survival in the context of acute infection.

Non-immune cells have important functions in pathogen defense. Understanding the earliest stages of host-fungal interactions is critical, and epithelial cells are some of the first cells to encounter conidia upon infection, along with AMs. In mucosal infection with *Candida albicans*, oral epithelial cells are known to respond to IL-17 signaling and produce anti-microbial peptides,^38,39^ and pulmonary epithelial cells use NF-κB signaling to promote innate lymphocyte antifungal responses during *Blastomyces* infection.^15^ However, the mechanisms by which epithelial cells communicate with neutrophils to regulate effector properties *in situ* remained undefined. Extensive *in vitro* and limited *in vivo* work has been done to investigate the function of pulmonary epithelial cells during *A. fumigatus* infection. Several groups have used epithelial cell lines (such as A549^40–42^ and HSAEC1-KT^43^) to demonstrate that *A. fumigatus* is endocytosed by epithelial cells in *in vitro* fungus-epithelial cell cultures. There is some evidence for endocytosis of *A. fumigatus* in murine models of IA during immunosuppression with corticosteroids that effectively eliminate leukocytes.^42^ While this may also occur in immunosuppressed patients, we propose that understanding the role of epithelial cells in immunocompetent settings will allow us to mimic their host-protective functions for therapeutic use in susceptible patient populations. In our model of immunocompetent murine infection with *A. fumigatus*, we found that a major function of epithelial cells is to produce GM-CSF. Additionally, GM-CSF levels were not impacted by loss of C-type lectin receptor signaling, but rather, SPC^+^ epithelial cells were responsive to other cytokine signals to increase GM-CSF production. We propose that an important function of epithelial cells during pulmonary infection may be to act as local signaling hubs to coordinate a site-specific anti-pathogen response. We posit that both the inducing signals for GM-CSF (i.e., IL-1 and IFN-λ) and the licensing function of GM-CSF on neutrophils exist in locally restricted areas of fungal infiltration and neutrophil accumulation, perhaps to limit immune-mediated damage only to areas of tissue with active infection, preserving tissue integrity and essential functions. This acute and spatially restricted process stands in contrast to the homeostatic production of GM-CSF by AECIIs, which must occur throughout the lung to support alveolar macrophages and thereby control surfactant levels that govern surface tension in the alveoli to prevent alveolar collapse.

In summary, we have described crosstalk between epithelial cells and neutrophils via GM-CSF as a key mechanism of defense against *A. fumigatus*, downstream of dual regulation by the cytokines IL-1 and IFN-λ. This study uncovers the critical, early, and spatially coordinated role of epithelial cells in promoting neutrophil-dependent protection against inhaled fungi, in addition to identifying the contribution of type III IFN signaling in early GM-CSF production in the lung. Future work may define the molecular and physical links between *A. fumigatus* and IL-1 and IFN-λ, the precise regulation of *A. fumigatus* killing by neutrophils downstream of GM-CSF signaling, and broader functions for both GM-CSF on other target cells and for epithelial cells as signaling hubs for orchestration of the pulmonary anti-*A. fumigatus* immune response.

## MATERIALS AND METHODS

### Mice

C57BL/6 mice (stock # 000664), *Csf2*^−/−^ mice (stock # 026812), *Sftpc*-Cre^ERT2^ mice (stock # 028054), *Il1r1*^−/−^ mice (stock # 003245), *Myd88*^−/−^ mice (stock # 009088), and *Ifnar1^−/−^* mice (stock # 032045-JAX) were purchased from The Jackson Laboratory. CD45.1^+^ mice (stock # 564) were purchased from Charles River Laboratories. *Il1r1*-flox (Jackson Laboratory stock # 028398) mice were provided by Dr. Iliyan Iliev (Weill Cornell Medicine). FROG mice^18^ (*Csf2* reporter), *Csf2*-floxed mice, and *Csf2rb*-floxed mice^26^ were provided by Dr. Burkhard Becher (University of Zurich). *Csf2*-floxed mice are described for the first time in this study and were generated at LTK Zürich (2018) using genetically modified JM8A3.N1 C57BL/6 stem cells from EUCOMM. ES cells were injected into C57BL/6 blastocysts and the resulting chimeric mice were bred with C57BL/6 to achieve germ line transmission and generate the knockout-first allele. The L1L2_GT0_LF2A_LacZ_BetactP_neo cassette was inserted by targeted mutation and subsequently the knockout-first mice were crossed with the flp delete cre resulting in a conditional ready allele (Fig. S5A). *Card9*^−/−^ mice^44^ were provided by Dr. Xin Lin (Tsinghua University). *Clec7a*^−/−^ mice^45^ were provided by Dr. Yoichiro Iwakura (Tokyo University of Science). *Fcer1g*^−/−^ mice were provided by Dr. Jessica Hamerman (Benaroya Research Institute). *Clec7a*^−/−^ *Fcer1g*^−/−^ mice have been previously described.^46^ *Mrp8*-Cre mice (Jackson Laboratory stock # 021614) were provided by Dr. Sunny Shin (University of Pennsylvania).

*Stat1*^−/−^ mice^22^ were provided by Dr. Joseph Sun (Memorial Sloan Kettering Cancer Center). *Ifnlr1*^−/−^ mice were provided by Dr. Sergei Kotenko (Rutgers University). *Ifnl2/3*^−/−^ mice were generated by the Rivera and Kotenko labs (Rutgers University)^47^. The mouse genome contains two functional IFN-λ genes, the *Ifnl2* and *Ifnl3* genes, which are juxtaposed in a head-to-head orientation on chromosome 7. Guide RNAs (gRNAs) were designed to target sequences flanking these genes, and CRISPR/Cas9 was used to generate mice lacking both IFN-λ genes, to create an *Ifnl2/3* knock-out (KO) strain. The selected *Ifnl2/3* KO strain contains a 19,803 base pair deletion (from 28,506,772 to 28,526,525 nucleotide positions; NCBI GRCm38.p4) that removes the entire *Ifnl2* and *Ifnl3* genes including their promoters and 3’UTR and replaces these with the ATAACTTCGTATAGCATA sequence.

All non-chimeric mice used in this study were 8-12 weeks old. Chimeric mice were 12-16 weeks old at time of infection. Within experiments, mice were age- and sex-matched. Experiments were performed with both male and female mice. Mouse strains, except for *Il1a/b*^−/−^ mice (NIH) and *Ifnl2/3*^−/−^ and *Ifnlr1*^−/−^ mice (Rutgers University), were bred and housed in the Research Animal Resource Center at MSKCC in individual ventilated cages under specific-pathogen free conditions. Animal experiments were conducted with approval of the MSKCC (protocol 13-07-008) or Rutgers University Institutional Animal Care and Use Committee (PROTO201900145) or following the recommendations in the Guide for the Care and Use of Laboratory Animals of the National Institutes of Health, under the auspices of protocol LCIM14E approved by the Animal Care and Use Committee of the NIAID. Animal studies complied with all applicable provisions established by the Animal Welfare Act and the Public Health Services Policy on the Humane Care and Use of Laboratory Animals.

### Aspergillus fumigatus *strains and murine infection model*

*Aspergillus fumigatus* strain CEA10 (isolate provided by Robert Cramer, Dartmouth University) was used for all experiments. For experiments to analyze fungal uptake and killing using FLARE conidia, the CEA10-mRFP strain was used.^48^ *A. fumigatus* conidia were grown on glucose minimal medium slants for 4-7 days at 37°C prior to harvesting in PBS + 0.025% Tween-20 for experimental use. For FLARE experiments, 7.5 x 10^8^ conidia were incubated in 10 μg/mL EZ-Link™ Sulfo-NHS-LC-Biotin (ThermoFisher) in 1 mL of 50 mM NaHCO_3_ buffer (pH 8.3) for 1-2 hrs at room temperature (or 16 hours at 4°C), washed with 1 mL of Tris-HCl (pH 8) buffer, incubated with 20 μg/mL Streptavidin, Alexa Fluor™ 633 conjugate (Molecular Probes) in PBS for 45 mins at room temperature, and resuspended in PBS + 0.025% Tween-20. To generate swollen heat-killed conidia, 5 x 10^6^/mL resting conidia were incubated for 16 hours in RPMI-1640 and 0.5 μg/mL voriconazole and then put in a heat block set to 100°C for 30 minutes. Mice were lightly anesthetized by isoflurane inhalation and 3-6 x 10^7^ *A. fumigatus* conidia were instilled via the intratracheal route in 50 μL of PBS + 0.025% Tween-20.

### Generation of bone marrow chimera mice

Recipient mice were lethally irradiated (900 cGy) and reconstituted with 2-5 x 10^6^ donor bone marrow cells. After transplantation, recipient mice received 400 μg/mL enrofloxacin in drinking water for 3 weeks to prevent bacterial infections. Mice were rested for 6-8 weeks prior to experimental use. 4 weeks post bone marrow transplant, chimerism was validated by bleeding mice from the tail, staining blood cells with antibodies against CD45.1 and CD45.2, and analyzing by flow cytometry. Samples from chimeric mice routinely stained >95% positive for the donor marker and <5% positive for the recipient marker.

### Tamoxifen inducible gene deletion

*Sftpc*-Cre^ERT2^ mice are engineered to have a tamoxifen-inducible Cre under the control of the *Sftpc* promoter.^49^ To delete *Il1r1* or *Csf2* in these mice by crossing to *Il1r1*-floxed or *Csf2*-floxed mice, 100 μL of a 40 mg/mL tamoxifen solution in corn oil was administered via the intraperitoneal route once per day on 4 consecutive days (4 mg per dose). Mice were rested for 10-12 days following the final injection before experimental use. In *Il1r1*^ΔSPC^ experiments, littermate control mice (Cre-negative) were also injected with tamoxifen to control for any tamoxifen-associated toxicity. In some *Csf2*^ΔSPC^ experiments, some Cre-positive mice were injected with vehicle (corn oil) alone as controls due to low numbers of Cre-negative mice.

### Preparation of mouse lungs for ELISA

Mouse lungs were dissected and put into 2 mL of PBS. Lungs were homogenized with a PowerGen 125 homogenizer (Fisher Scientific) for 10-15 seconds. Samples were centrifuged at 1500 rpm for 5 minutes and supernatants were pipetted into a deep-well 96 well plate for long term storage at −80°C.

### Quantification of fungal burden

To measure colony-forming units (CFU) in the lungs of infected mice, lungs were dissected and homogenized with a PowerGen 125 homogenizer (Fisher Scientific) for 10-15 seconds in 2 mL of PBS. 10 μL was removed and diluted for plating onto Sabourand dextrose agar plates. Plates were incubated for 48 hours at 37°C and CFU were enumerated by counting. To quantify germination, mouse lungs were perfused with PBS through the right ventricle, collected into histology cassettes, stored in 4% paraformaldehyde for 16 hours, then placed in ethanol. Samples were embedded in paraffin, sectioned, and stained with Gomori methenamine silver (GMS) to identify fungal organisms. Slides were scanned using a Pannoramic Digital Slide Scanner (3DHISTECH, Budapest, Hungary) using a 20x/0.8NA objective. Images were color deconvolved to separate the ubiquitous green stain from the black fungal stain (GMS) and annotated in CaseViewer 2.4 (3DHISTECH). The fungi were segmented (outlined) and the longest diameter of the bounding box that surrounded each fungal organism (Feret’s diameter) was calculated for each fungal organism. Organisms with a Feret diameter >6 μm were classified as germinating. The number of germinating organisms was divided by the total number of organisms and multiplied by 100 to calculate the germination percentage.

### Flow cytometry

For analysis of immune cells, single cell suspensions of mouse lungs were generated by putting lungs in gentle MACS™ C tubes and mechanically homogenizing in 5 ml PBS using a gentle MACS™ Octo Dissociator (Miltenyi Biotec) in the absence of enzymes, then filtered through 100 μm filters. For analysis of lung structural cells, single cell suspensions were generated as in^50^, with minor modifications. Briefly, mice were euthanized by pentobarbital injection, dissected to reveal the trachea and chest cavity, and a tracheal cannula was inserted. Lungs were perfused with 5 mL of PBS via injection through the right ventricle. 1 mL of Dispase was pushed through the right ventricle to coat the lung endothelium and 0.7 mL of Dispase was pushed through the tracheal cannula to coat the lung epithelium, followed by 0.5 mL 1% low-melting point agarose through the tracheal cannula to prevent leakage of the enzymes. The lungs were then covered in ice to semi-solidify the agarose before removal into 5 mL of cold PBS. Lungs were mechanically digested using scissors into pieces <2 mm^2^ and put in 5 mL of digestion buffer (PBS + 0.5% BSA + 5 mg/mL Collagenase I + 100 μg/mL DNase I) in 15 mL conical tubes on a rotator at 37°C for 45 minutes. A 1 mL pipette was used to disrupt any further clumps of cells by pipetting 50 times. Lung suspensions were then filtered through 100 μm filters. For immune and structural cell analysis, red blood cells were lysed using RBC lysis buffer (Tonbo Biosciences), cells were blocked with anti-CD16/CD32, stained with fluorophore-conjugated antibodies, and analyzed on a Beckman Coulter Cytoflex LX. Single color controls for compensation were generated using lung cells or OneComp eBeads™ Compensation Beads. Experiments were analyzed with FlowJo version 10.8.1. Dead cells were excluded with DAPI or eBioscience™ Fixable Viability Dye eFluor™ 506 (ThermoFisher). Gating strategies for all cell subsets described in the manuscript are provided in the supplementary figures, except for the lymphocyte subsets in Fig. S3E-S3H. These are defined as follows: γδ T cells are CD45^+^ CD11b^−^ CD3^+^ TCRδ^+^, ILCs are CD45^+^ CD3^−^ CD90.2^+^, αβ T cells are CD45^+^ CD11b^−^ CD3^+^ TCRβ^+^, and NK cells are CD45^+^ CD3^−^ CD11b^+^ NK1.1^+^.

### Preparation of lung samples for sorting

Single cell lung suspensions were prepared for sorting using the protocol for structural cell enrichment for flow cytometry. Sorting was performed on a BD FACSAria™ III using a 100 μm nozzle in the Flow Cytometry Core Facility at MSKCC. Cells were collected into 1 mL ice cold FBS.

### Quantitative PCR

RNA was isolated using TRIzol™ LS Reagent (ThermoFisher) or RNeasy Mini Kit (Qiagen) and following the manufacturer protocol. For qPCR, cDNA was generated from RNA by using the High-Capacity RNA-to-cDNA™ Kit (Applied Biosystems) and following the manufacturer protocol. qPCR was performed on a StepOnePlus Real Time PCR System (Applied Biosystems) using TaqMan Fast Advanced Master Mix and TaqMan Gene Expression Assays (ThermoFisher Scientific) for *Csf2*, *Csf2rb*, *Sftpc*, and *Scgb1a1*, and results were normalized to ribosomal RNA quantities using TaqMan™ Ribosomal RNA Control Reagents (Applied Biosystems). For quantification of *Il1r1* mRNA, PowerUp™ SYBR™ Green Master Mix (ThermoFisher) was used and results were normalized to *Gapdh* mRNA.

### Analysis of fungal uptake and killing

Mice were infected with FLARE conidia and lungs were harvested and prepared for flow cytometry. Uptake refers to the proportion of fungus-engaged cells, i.e., the fraction of a given cell subset that are AF633^+^ (the sum of RFP^+^ AF633^+^ and RFP^−^ AF633^+^ cells). Viability refers to the proportion of fungus-engaged cells that contain live conidia, i.e., RFP^+^ AF633^+^ cells divided by the sum of RFP^+^ AF633^+^ and RFP^−^ AF633^+^ cells.

### Immunofluorescence imaging

Lungs were perfused with 4% paraformaldehyde or PLP solution (Cold Spring Harbor Protocols^51^) and individual lung lobes were dissected and placed in 2% paraformaldehyde on a rotator for 16 hours at 4°C. Lungs were washed with PBS and put in 30% sucrose in PBS for 24-48 hours at 4°C. Lung lobes were embedded in OCT, frozen on dry ice, and transferred to - 80°C for long term storage. Samples were cryo-sectioned at 10-20 μm onto Superfrost™ Plus Microscope Slides for storage at −20°C. A hydrophobic barrier was drawn on thawed slides using an ImmEdge® Hydrophobic Barrier PAP Pen, the OCT was dissolved with PBS. Samples were blocked for 1 hour at room temperature using 10% donkey serum diluted in Animal-Free Blocker® and Diluent, R.T.U. (Vector Laboratories). Primary staining was done at 4°C overnight, secondary staining at room temperature for 2 hours, and DAPI staining at room temperature for 10-20 mins. For samples infected with AF633-labeled *A. fumigatus*, the AF633 fluorescence emission was strong enough to be detected without amplification, and slides were stained with GFP (AF488), pro-SPC (AF594), and DAPI. For samples infected with unlabeled *A. fumigatus* to be analyzed for distances to neutrophils, slides were stained with GFP (AF488), pro-SPC (AF594), Ly6G (AF647), and DAPI. Samples were imaged using a Leica SP8 using a 40x/1.1NA objective (Fig. 4G, 5B & D) or a Pannoramic Digital Slide Scanner (3DHISTECH, Budapest, Hungary) using a 40x/0.95NA objective (Fig. 5C & E, Fig. S4A-M).

### *Quantification of distances from GM-CSF-producing cells to* A. fumigatus *and neutrophil*

In ImageJ, cells were segmented (outlined) based on DAPI staining. DAPI^+^ objects were classified as positive or negative for each of the other markers, based on the presence or absence of overlap with positive pixels from each respective fluorophore. The minimum distance from each SPC^+^ (AF594) object to either *A. fumigatus* (AF633) or neutrophils (AF647) was determined by generating a Euclidian distance map to the positive pixels for *A. fumigatus* (AF633) or to Ly6G (AF647). Each AF594^+^ (pro-SPC) cell’s distance was calculated as the mean of all the pixel’s distances that comprise the cell. SPC^+^ cells were categorized as GM- CSF^+^ or GM-CSF^−^ based on GFP fluorescence, and distance measurements in the two groups were compared.

## Key resources table

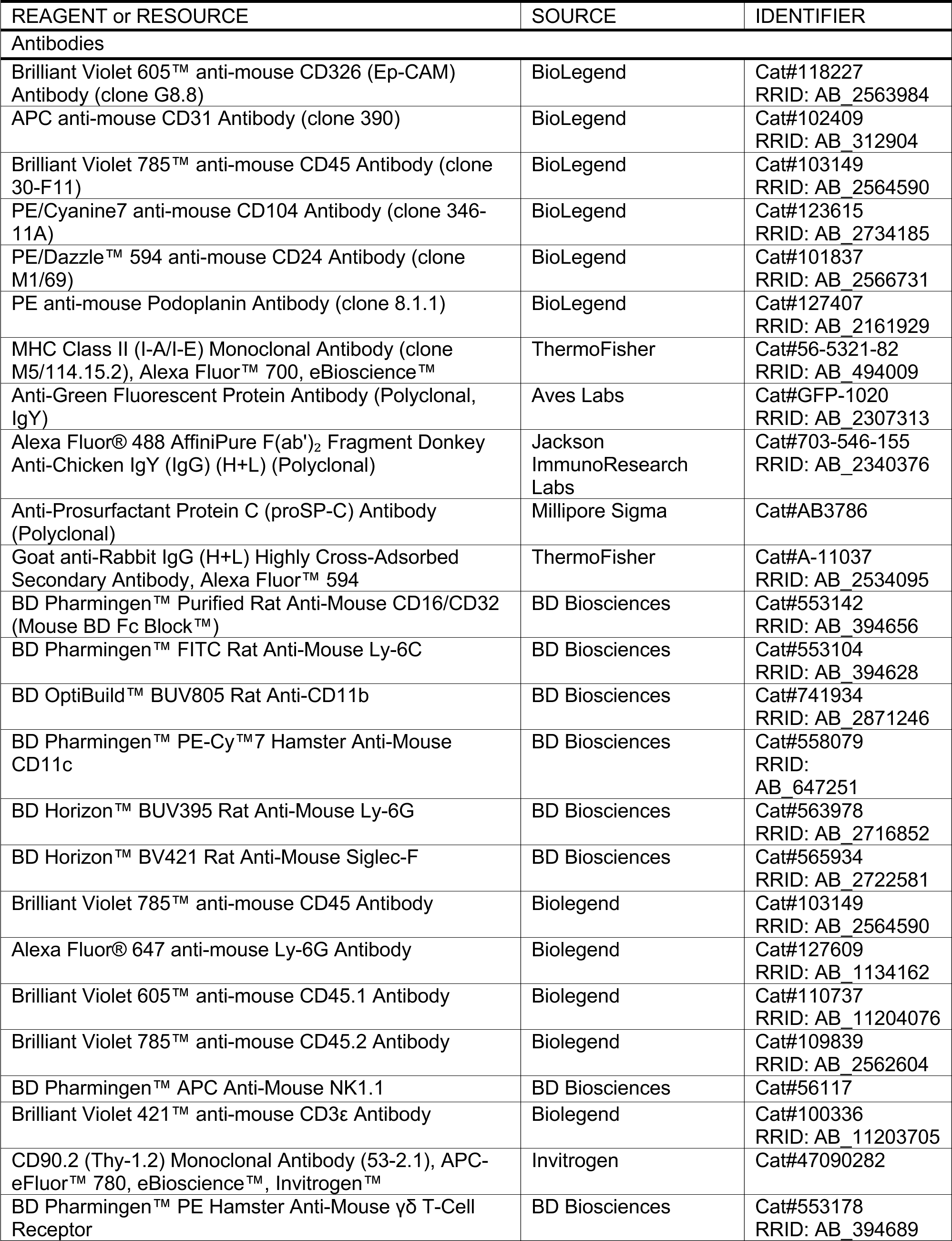
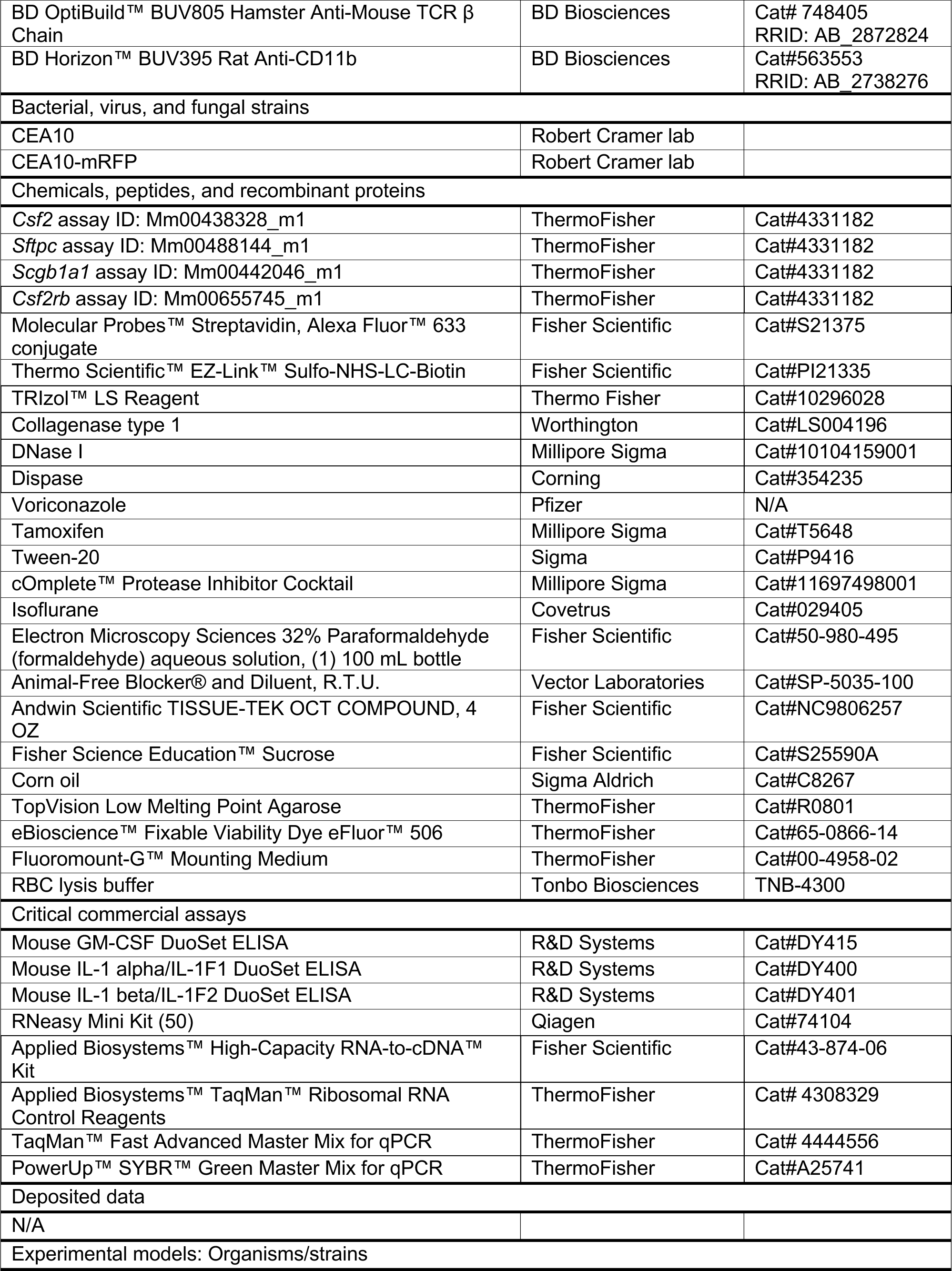
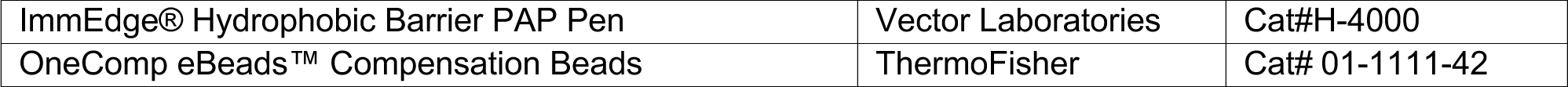

## ACKNOWLEDGEMENTS

We thank all members of the Hohl lab for insightful discussions. We thank Robert Cramer (Dartmouth College) for sharing the *A. fumigatus* strains used in this work. We thank Sunny Shin (U. Pennsylvania) for sharing *Csf2rb*^ΔNeuts^ mice and for helpful discussion. We thank Joseph Sun (MSKCC) for sharing *Stat1*^−/−^ mice, Iliyan Iliev (Weill Cornell Medicine) for sharing *Il1r1*-flox mice, Jessica Hamerman (Benaroya Research Institute) for sharing *Fcer1g*^−/−^ mice, Xin Lin (Tsinghua University) for sharing *Clec9a*^−/−^ mice, Katrin Mayer-Barber (NIAID/NIH) for sharing *Il1a/b*^−/−^ mice, and Sergei Kotenko (Rutgers University) for sharing *Ifnlr1*^−/−^ mice. We acknowledge the MSKCC Molecular Cytology Core Facility (Murray Tipping, Michael Galiano, Ning Fan) and Deeksha Deep (MSKCC) for assistance with imaging techniques and the MSKCC Flow Cytometry Core Facility for assistance with cell sorting. We gratefully acknowledge technical assistance from Mergim Gjonbalaj and Audrey Billips. The graphical abstract, Fig. 5A, Fig. 6C, and Fig. S5A were created with BioRender.com. These studies were supported by NIH grants T32 AI134632 (Weill Cornell Graduate School, PI: Sabine Ehrt & Ming Li), F31 AI167511 (KAMM), F31 AI161996 (MAA), K99/R00 AI141622 (JVD), P30 CA008748 (MSKCC, PI: Selwyn Vickers), R01 AI169769-01 (AR), R01 AI169770-01A1 (AR), R37 AI093808 (TMH), R01 AI139632 (TMH), and the Division of Intramural Research of the NIAID/NIH (ZIA AI0011175 to MSL). These studies were also supported by the University of Zurich Forschungskredit Postdoctoral Fellowship K-41302-10-01 (DDF), the Swiss Multiple Sclerosis Society (DDF), the European Research Council (ERC) under the European Union’s Horizon 2020 Research and Innovation Program Grant Agreement # 882424 (BB), the Swiss National Science Foundation # 310030_188450 (BB), and the Burroughs Wellcome Fund award # BWF-PATH 1015502 (AR). The funders had no role in study design, data collection and analysis, decision to publish, or preparation of manuscript.

## AUTHOR CONTRIBUTIONS

Conceptualization: K.A.M.M. and T.M.H.

Methodology: K.A.M.M. and E.R.

Formal Analysis: K.A.M.M. and E.R.

Investigation: K.A.M.M., V.E., J.V.D., M.A.A., Y.G., K.A.M.

Resources: F.R., S.T., D.D.F., A.R., B.B.

Writing – Original Draft: K.A.M.M., F.R., E.R., V.E., A.R., B.B., T.M.H.

Writing – Reviewing and Editing: K.A.M.M. and T.M.H.

Supervision: M.S.L., A.R., B.B., T.M.H.

Funding Acquisition: K.A.M.M., M.A.A., D.D.F., M.S.L., A.R., B.B., T.M.H.

---

**Fig. S1, related to Fig. 1.**
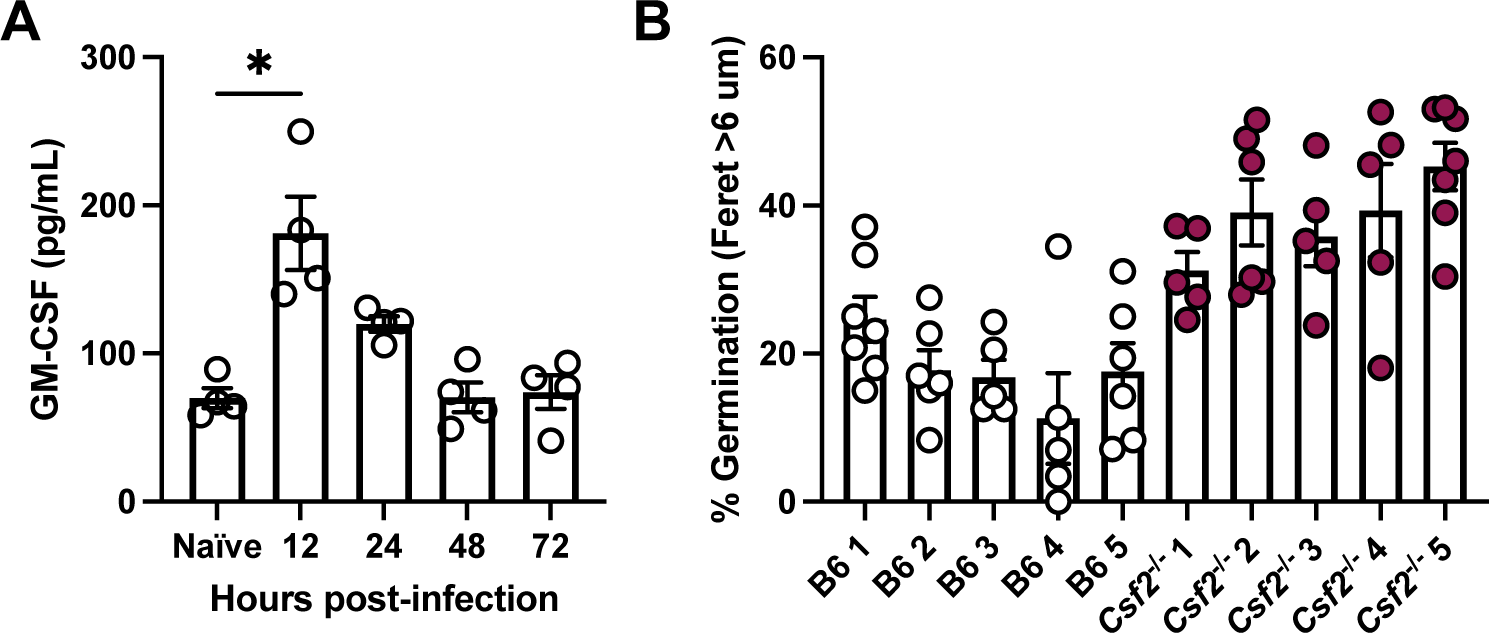
GM-CSF is induced during *A. fumigatus* infection and restricts conidial germination. (A) Representative experiment of GM-CSF levels measured by ELISA in naïve and *A.f.* infected (3 x 10^7^) B6 mice. Each symbol is one mouse. Significance determined by Kruskal-Wallis test. (B) Representative experiment of percentage of germinating conidia quantified from GMS staining of lung sections. Each bar is one mouse, and each dot is one annotated image area analyzed. (A & B) Data is displayed as mean ± SEM.

**Fig. S2, related to Fig. 2.**
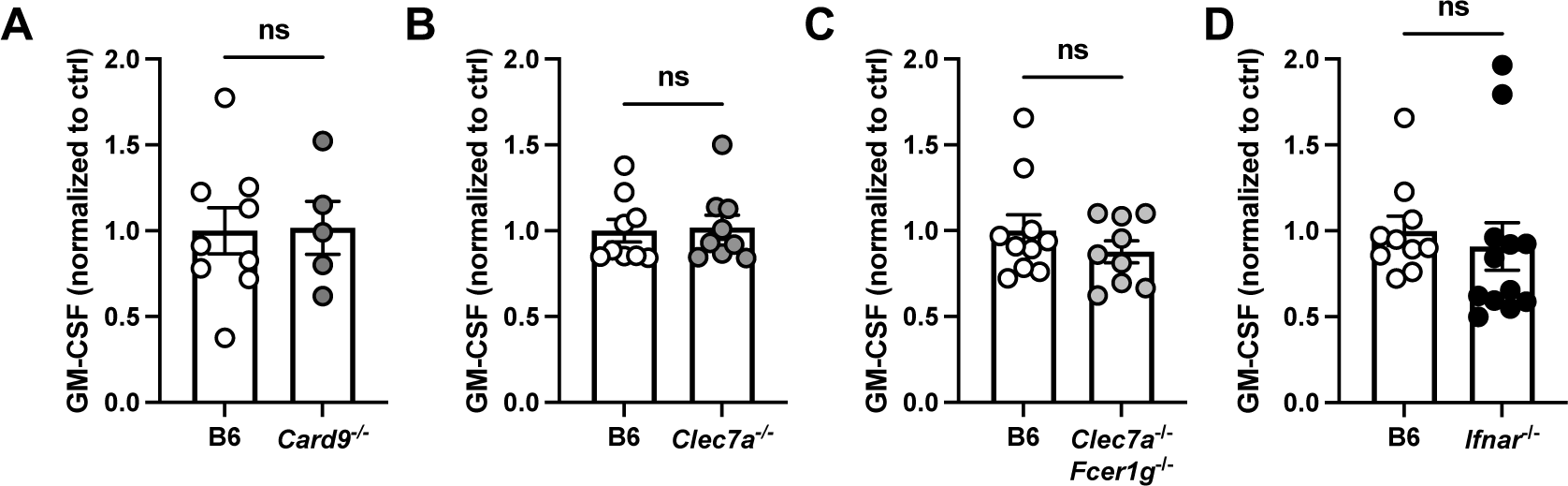
C-type lectin receptor signaling and type I IFN signaling are dispensable for GM-CSF production during *A. fumigatus* infection. (A-D) GM-CSF levels measured by ELISA in B6 and (A) *Card9*^−/−^, (B) *Clec7a*^−/−^, (C) *Clec7a*^−/−^ *Fcer1g*^−/−^ and (D) *Ifnar*^−/−^ mice. 2-3 experiments pooled; each symbol represents one mouse and data is displayed as mean ± SEM. Significance determined by Mann-Whitney test.

**Fig. S3, related to Fig. 4.**
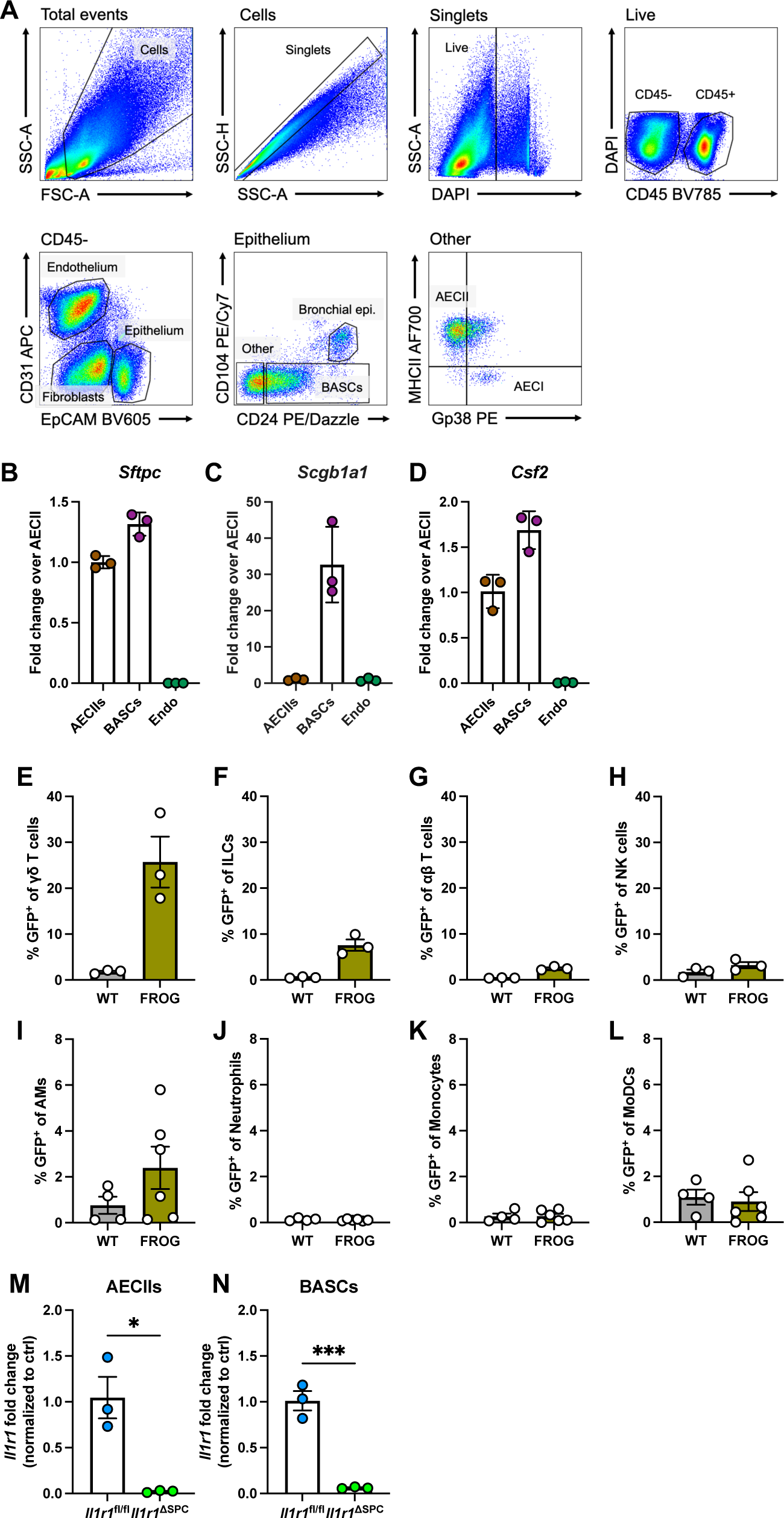
Validation of murine models used to identify GM-CSF-producing cell types. (A) Gating strategy for non-hematopoietic cells in the mouse lung. (B-D) Expression of (B) *Sftpc*, (C) *Scgb1a1*, and (D) *Csf2* measured by qPCR in FACS sorted AECIIs, BASCs, and endothelial cells from lungs of uninfected mice. (E-L) Percentage of GFP^+^ cells: (E) γδ T cells, (F) ILCs, (G) αβ T cells, (H) NK cells, (I) alveolar macrophages, (J) neutrophils, (K) monocytes, (L) monocyte-derived DCs. Analysis performed by flow cytometry 24 hpi. (M-N) *Il1r1* transcript levels in FACS sorted (M) AECIIs or (N) BASCs from uninfected *Il1r1*^fl/fl^ (Cre-negative) and *Il1r1*^ΔSPC^ mice. Significance calculated by Mann-Whitney test. (B-N) Each symbol represents one mouse.

**Fig. S4, related to Fig. 5.**
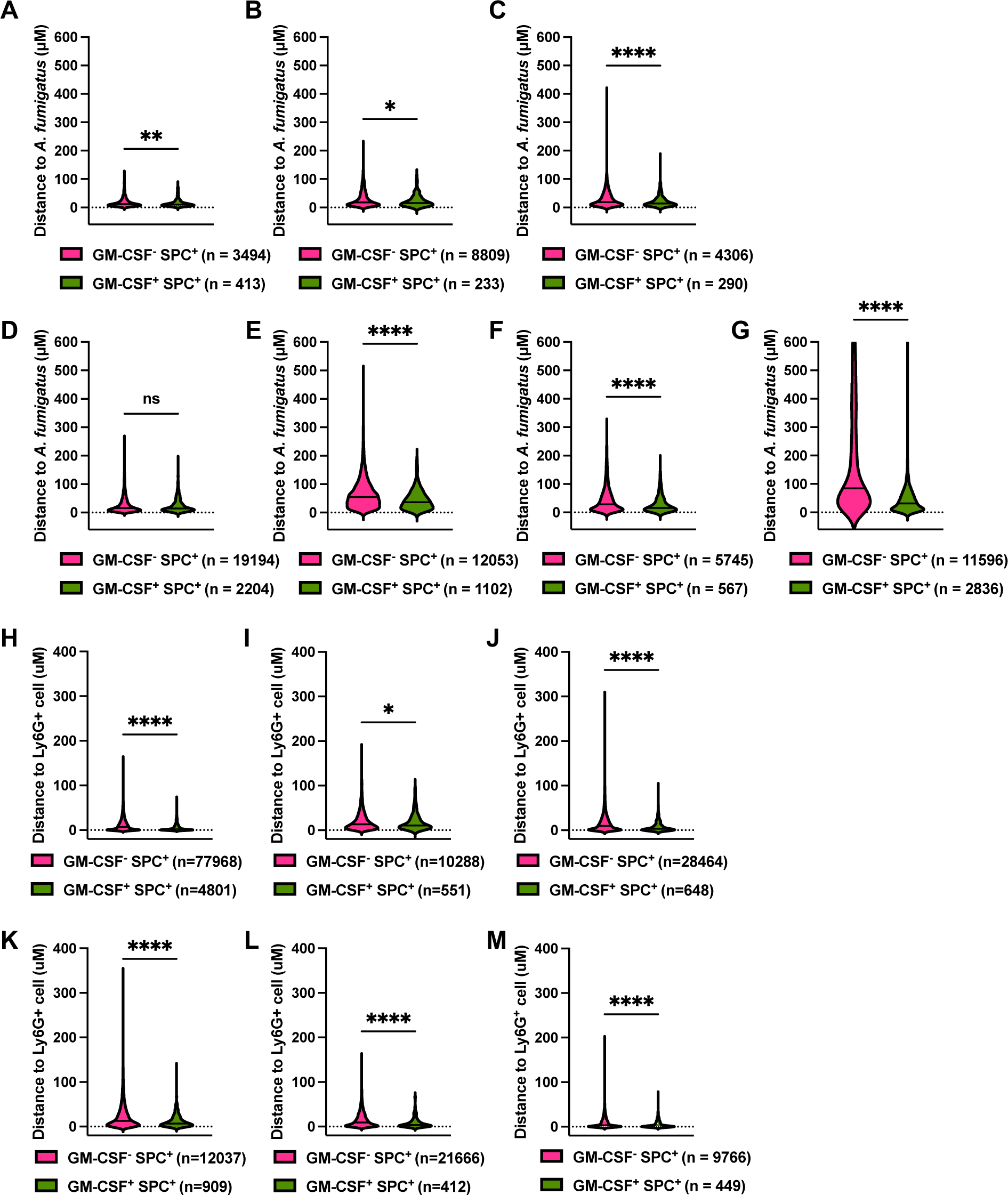
Distance of GM-CSF-producer and non-producer SPC^+^ epithelial cells to *A. fumigatus* and to neutrophils in individual biological replicates. Each chart represents immunofluorescence data from one mouse and cell number in each group is indicated below the graph. (A-G) Violin plot displaying distance to nearest AF633^+^ *A. fumigatus* from indicated cell type. All mice are compiled in Fig. 5C. (H-M) Violin plot displaying distance to nearest Ly6G^+^ cell from indicated cell type. All mice are compiled in Fig. 5E.

**Fig. S5, related to Fig. 6.**
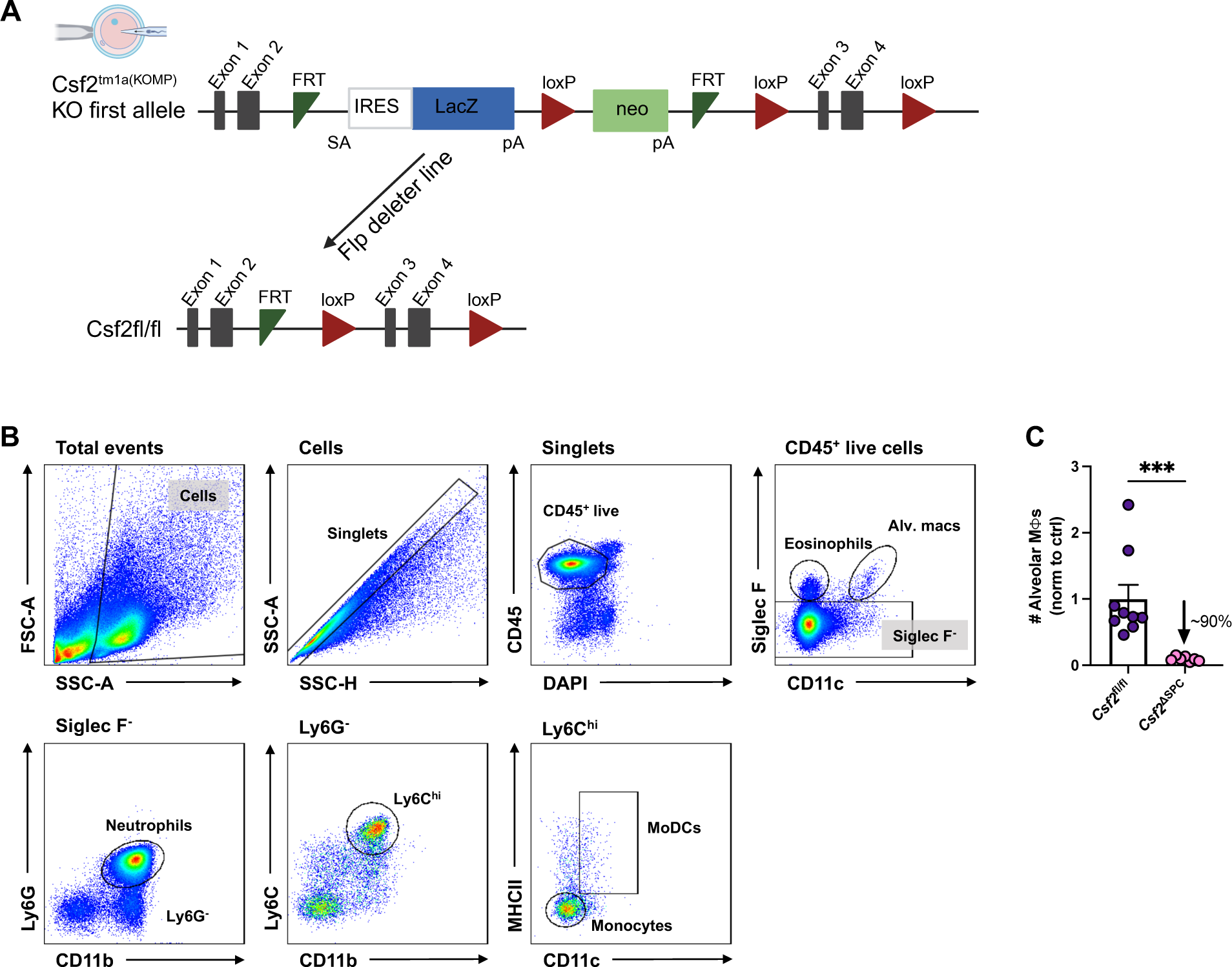
Generation of *Csf2*-floxed mice and analysis of lung immune cells by flow cytometry. (A) Generation of *Csf2*-floxed mice. (B) Gating strategy for myeloid cells in the *A. fumigatus*-infected mouse lung. (C) Number of alveolar macrophages in control and *Csf2*^Δ*SPC*^ mouse lungs 24 hpi. Data are from two pooled experiments. Each symbol represents one mouse. Data were analyzed by Mann-Whitney test.

**Fig. S6, related to Fig. 7.**
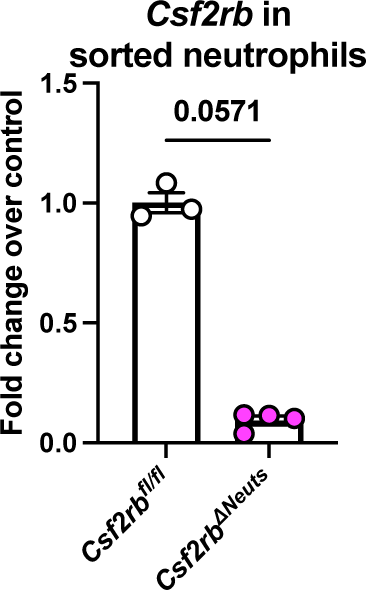
Validation of the *Csf2rb*^ΔNeuts^ mouse strain. *Csf2rb* transcript levels measured by qPCR in FACS sorted neutrophils from *Csf2rb*^fl/fl^ and *Csf2rb*^ΔNeuts^ mice 24 hpi. Significance calculated by Mann-Whitney test.

